# Pan-cancer analysis in the real-world setting uncovers immunogenomic drivers of acquired resistance post-immunotherapy

**DOI:** 10.1101/2025.07.04.663141

**Authors:** Mohamed Reda Keddar, Sebastian Carrasco Pro, Roy Rabbie, Zeynep Kalender Atak, Ana Camelo Stewart, Scott A. Hammond, Douglas C. Palmer, Ross Stewart, Kathleen Burke, Ben Sidders, Jessica Davies, Jonathan Dry, Inigo Martincorena, Sajan Khosla, Adam J. Schoenfeld, Martin L. Miller

## Abstract

Immune checkpoint blockade (ICB) has revolutionised cancer therapy, yet resistance — both primary and acquired — remains a significant obstacle, affecting the majority of patients. Here, we leveraged a large-scale, real-world clinicogenomic dataset to systematically explore the molecular underpinnings of ICB resistance in the post-progression setting. Analysing over 5,000 pan-cancer patients with clinical and pre-/post-treatment genomic and transcriptomic data, we identify distinct immunogenomic drivers of acquired vs. primary ICB resistance. Post-ICB progression, acquired resistance showed extended survival compared to primary resistance across all cancer types. The acquired resistance clinical phenotype was paralleled by a universally immune-inflamed, albeit dysfunctional, tumour microenvironment (TME) at the onset of acquired resistance, with sustained or ICB-induced inflammatory and interferon responses. We confirm previously described mechanisms of acquired resistance, including *B2M* loss-of-function (LoF) in non-small cell lung cancer (NSCLC), and identify novel potential mediators, including LoF of *TGFBR2* in NSCLC, *CYLD* in head and neck cancer, and *RUNX1* in triple negative breast cancer. Further supporting their involvement in resistance, these acquired ICB alterations associated with immune-escaped TMEs, characterised by active immunomodulatory oncogenic signalling, hyperproliferation and invasiveness, or altered tumour metabolism. These findings emphasise the heterogeneity of molecular drivers of acquired resistance to ICB within and across cancers, and highlight the potential for personalised therapeutic interventions post-progression to improve patient outcomes.

## INTRODUCTION

Immune checkpoint blockade (ICB) has deeply transformed the treatment landscape of cancer, increasingly becoming established as the standard of care (SoC) across many solid malignancies^1,2^. While a subset of patients can perform exceptionally well on ICB, 40-90% of cases derive no benefit from treatment (primary resistance) and up to 70% of initial responders eventually relapse and progress, yielding ∼5-25% of patients with acquired resistance^3,4^. Significant translational efforts have been put towards understanding molecular correlates of response and resistance to ICB, focusing primarily on analysing pre-treatment tumour biopsies from responders and non-responders. Such efforts contributed to the development of robust response biomarkers, some of which are approved by the US Food and Drug Administration (FDA) for patient selection and stratification. These include high tumour mutational burden (TMB), microsatellite instability or mismatch repair deficiency (MSI/dMMR), or high protein expression of the programmed cell death ligand 1 (PD-L1) immune checkpoint^4,5^.

Despite these efforts, our understanding of which molecular features emerge as patients progress on ICB remains very limited, particularly in the context of acquired resistance. This is notably due to challenges in accessing post-treatment clinical samples, as well as the lack of a clear and standardised definition of acquired resistance^3,6,7^. So far, efforts to address this gap, including ours^8^, focused on analysing (un)paired post-ICB tumour biopsies from mostly non-small cell lung cancer (NSCLC) or melanoma patients, uncovering several mechanisms underlying acquired ICB resistance. These mechanisms include genetic defects in antigen processing and presentation machinery (APM) and/or interferon (IFN) signalling^8–13^, chronic IFN gamma (IFNγ) signalling^8^, cancer driver alterations promoting immune exclusion^14,15^, or upregulation of alternative immune checkpoints following treatment^16–18^. The diversity of these mechanisms underscores the heterogeneous nature of acquired resistance to ICB and suggests that additional mediators may have been overlooked due to limited sample sizes. Moreover, the extent to which these mechanisms are shared in other cancer types, in post-ICB primary resistance, or in acquired resistance to non-ICB regimens remains underexplored. Addressing these questions is crucial to direct therapeutic interventions post-ICB resistance towards more personalised approaches.

The establishment of ICB as part of routine care comes with an ever-increasing availability of clinical samples across various treatment settings, including the post-progression setting. Therefore, clinicogenomic real-world data (RWD) provides an exciting opportunity to dissect the molecular drivers of resistance as it emerges across multiple cancer types. Here, we leverage a real-world dataset from Tempus AI (tempus.com) to systematically analyse clinical and pre-/post-treatment multi-omic data from >5,000 patients with cancer. We uncover molecular determinants of acquired resistance across various disease settings, providing the largest-to-date and first cross-indication and multi-modal comparison of acquired vs. primary ICB resistance.

## RESULTS

### Building a pan-cancer multi-modal ICB cohort from real-world data

We leveraged Tempus AI’s US-based and de-identified clinicogenomic database to build a pan-cancer and multi-modal real-world ICB (rwICB) cohort with follow-up until August 2023. Available data includes treatment history and outcomes from patient electronic health records (EHR), panel DNA sequencing (DNA-seq), full transcriptome RNA-seq, and associated sample and clinical metadata (Supplementary Figure 1A). Following a systematic informatics approach, we started from 22,952 patients spanning ten cancer types and applied a set of inclusion and exclusion criteria to build rwICB (Supplementary Figure 1B, Methods). We considered patients who had 1) received at least one line of ICB (monotherapy or in combination with other drugs) in their treatment journey, 2) complete treatment information, 3) treatment outcomes, and 4) omic data (DNA and/or RNA) for analysis. All disease stages were considered. This allowed us to identify 3,559 patients across NSCLC (n=2,689), head and neck squamous cell cancer (HNC, n=516), and triple-negative breast cancer (TNBC, n=354) who received ICB between 2016 and 2023. To contrast our findings with other treatment modalities, we built a homologous and non-overlapping control cohort of 1,894 non-ICB-treated patients, consisting of chemotherapy for all three cancer types, as well as tyrosine kinase inhibitors (TKI) for NSCLC (Supplementary Figure 1B). The baseline patient characteristics of the ICB and non-ICB cohorts are summarised in Table 1.

**Table 1.** Baseline patient characteristics of the ICB and non-ICB multi-modal cohorts. For age and treatment start year, both median and range (min-max) are reported. For TNBCs in the chemotherapy cohort, 51/503 (10%) patients received chemotherapy between 1991-2015. All other ICB and non-ICB patients received treatment between 2016-2023. For age, the number of patients missing age information (N/A) is reported in brackets. For follow-up, the number of patients missing or with inconsistent last known follow-up date and/or censoring information is reported in brackets (N/A). ‘anti-PDx mono’ designates anti-PD(L)1 administered as monotherapy, ‘anti-PDx + chemo’ designates anti-PD(L)1 administered concomitantly with chemotherapy, ‘Other ICB’ includes anti-PD1 combined with anti-CTLA-4, anti-PD(L)1 combined with other non-ICB drugs, or any anti-PD(L)1 part of a sequential drug combination. NSCLC = non-small cell lung cancer, HNC = head and neck squamous cell cancer, TNBC = triple-negative breast cancer, ECOG = Eastern Cooperative Oncology Group performance status, LoT = line of treatment, anti-PDx = anti-PD(L)1, mono = monotherapy, chemo = chemotherapy, TKI = tyrosine kinase inhibitors.

|  | ICB |  |  | Chemotherapy |  |  | TKI |
| --- | --- | --- | --- | --- | --- | --- | --- |
|  | NSCLC<br>(n=2689) | HNC<br>(n=516) | TNBC<br>(n=354) | NSCLC<br>(n=574) | HNC<br>(n=462) | TNBC<br>(n=503) | NSCLC<br>(n=355) |
| Median age at treatment start | 66<br>(21-83) | 62<br>(26-87) | 57<br>(29-84) | 66<br>(31-83, 3 N/A) | 60<br>(21-86) | 57<br>(25-84, 1 N/A) | 65<br>(30-87) |
| Median treatment start year | 2019<br>(2016-2023) | 2020<br>(2016-2022) | 2020<br>(2016-2022) | 2019<br>(2016-2022) | 2020<br>(2016-2022) | 2018<br>(1991-2022) | 2019<br>(2016-2022) |
| Median follow-up (months) | 18 (19 N/A) | 13 (4 N/A) | 16 | 19 (4 N/A) | 15 | 25 (1 N/A) | 26 (2 N/A) |
| **Gender** |  |  |  |  |  |  |  |
| Female | 1285 (47.8%) | 121 (23.4%) | 353 (99.7%) | 300 (52.3%) | 113 (24.5%) | 500 (99.4%) | 243 (68.5%) |
| Male | 1404 (52.2%) | 395 (76.6%) | 1 (0.3%) | 274 (47.7%) | 349 (75.5%) | 3 (0.6%) | 112 (31.5%) |
| **Ethnicity** |  |  |  |  |  |  |  |
| White | 1605 (59.7%) | 293 (56.8%) | 174 (49.2%) | 325 (56.6%) | 207 (44.8%) | 232 (46.1%) | 176 (49.6%) |
| Non-white | 472 (17.6%) | 69 (13.4%) | 84 (23.7%) | 87 (15.2%) | 70 (15.2%) | 122 (24.3%) | 93 (26.2%) |
| N/A | 612 (22.8%) | 154 (29.8%) | 96 (27.1%) | 162 (28.2%) | 185 (40%) | 149 (29.6%) | 86 (24.2%) |
| **Smoking history** |  |  |  |  |  |  |  |
| Never smoker | 144 (5.4%) | 62 (12%) | 98 (27.7%) | 39 (6.8%) | 22 (4.8%) | 46 (9.1%) | 72 (20.3%) |
| Ever smoker | 1053 (39.2%) | 139 (26.9%) | 49 (13.8%) | 182 (31.7%) | 95 (20.6%) | 53 (10.5%) | 45 (12.7%) |
| N/A | 1492 (55.5%) | 315 (61%) | 207 (58.5%) | 353 (61.5%) | 345 (74.7%) | 404 (80.3%) | 238 (67%) |
| **NSCLC histology** |  |  |  |  |  |  |  |
| Non-squamous | 1963 (73%) | N/A | N/A | 440 (76.7%) | N/A | N/A | 350 (98.6%) |
| Squamous | 726 (27%) | N/A | N/A | 134 (23.3%) | N/A | N/A | 5 (1.4%) |
| **ECOG** |  |  |  |  |  |  |  |
| 0-1 | 420 (15.6%) | 0 | 68 (19.2%) | 80 (13.9%) | 0 | 32 (6.4%) | 50 (14.1%) |
| ≥2 | 60 (2.2%) | 0 | 3 (0.8%) | 4 (0.7%) | 0 | 1 (0.2%) | 4 (1.1%) |
| N/A | 2209 (82.1%) | 516 (100%) | 283 (79.9%) | 490 (85.4%) | 462 (100%) | 470 (93.4%) | 301 (84.8%) |
| **Stage** |  |  |  |  |  |  |  |
| ≤Stage III | 658 (24.5%) | 25 (4.8%) | 39 (11%) | 357 (62.2%) | 66 (14.3%) | 244 (48.5%) | 16 (4.5%) |
| Stage IV | 1935 (72%) | 406 (78.7%) | 299 (84.5%) | 168 (29.3%) | 305 (66%) | 227 (45.1%) | 327 (92.1%) |
| N/A | 96 (3.6%) | 85 (16.5%) | 16 (4.5%) | 49 (8.5%) | 91 (19.7%) | 32 (6.4%) | 12 (3.4%) |
| **LoT** |  |  |  |  |  |  |  |
| 1<sup>st</sup> | 1448 (53.8%) | 186 (36%) | 89 (25.1%) | 574 (100%) | 462 (100%) | 503 (100%) | 355 (100%) |
| ≥2<sup>nd</sup> | 1241 (46.2%) | 330 (64%) | 265 (74.9%) | 0 | 0 | 0 | 0 |
| **ICB regimen** |  |  |  |  |  |  |  |
| anti-PDx mono | 720 (26.8%) | 263 (51%) | 28 (7.9%) | N/A | N/A | N/A | N/A |
| anti-PDx + chemo | 1331 (49.5%) | 181 (35.1%) | 265 (74.9%) | N/A | N/A | N/A | N/A |
| Other ICB | 638 (23.7%) | 72 (14%) | 61 (17.2%) | N/A | N/A | N/A | N/A |
| **TKI regimen** |  |  |  |  |  |  |  |
| EGFR inhibitors | N/A | N/A | N/A | N/A | N/A | N/A | 283 (79.7%) |
| ALK inhibitors | N/A | N/A | N/A | N/A | N/A | N/A | 61 (17.2%) |
| Other TKI | N/A | N/A | N/A | N/A | N/A | N/A | 11 (3.1%) |

We then used clinical follow-up and real-world outcomes from the Tempus database to annotate long-term response, acquired or primary resistance to treatment (Figure 1A). Long-term response was defined as documented Complete Response (CR), Partial Response (PR) or durable Stable Disease (SD), with no report of progression throughout a follow-up period lasting at least six months. Primary resistance was defined as evidence of progression via Progressive Disease (PD), report of disease recurrence or metastasis, or presence of a subsequent line of treatment (LoT), with no evidence of initial response. Lastly, acquired resistance was defined as evidence of initial response (CR/PR/SD) followed by evidence of progression^19,20^ (PD, recurrence/metastasis, or subsequent LoT, see Methods for detailed definitions). We implemented these definitions for both ICB– and non-ICB-treated patients to compare and contrast our findings across cancer types and treatment modalities without definition-related biases. In both the ICB and non-ICB cohorts, long-term response was mostly evidenced by report of CR/PR, while primary resistance consisted mainly of PD or report of tumour recurrence or metastasis in the patient EHR (Supplementary Figures 1C-D). Similarly, only a minority of acquired resistant patients had durable SD as their initial response or presence of a subsequent LoT as their evidence of progression (Supplementary Figure 1E).

**Figure 1.**
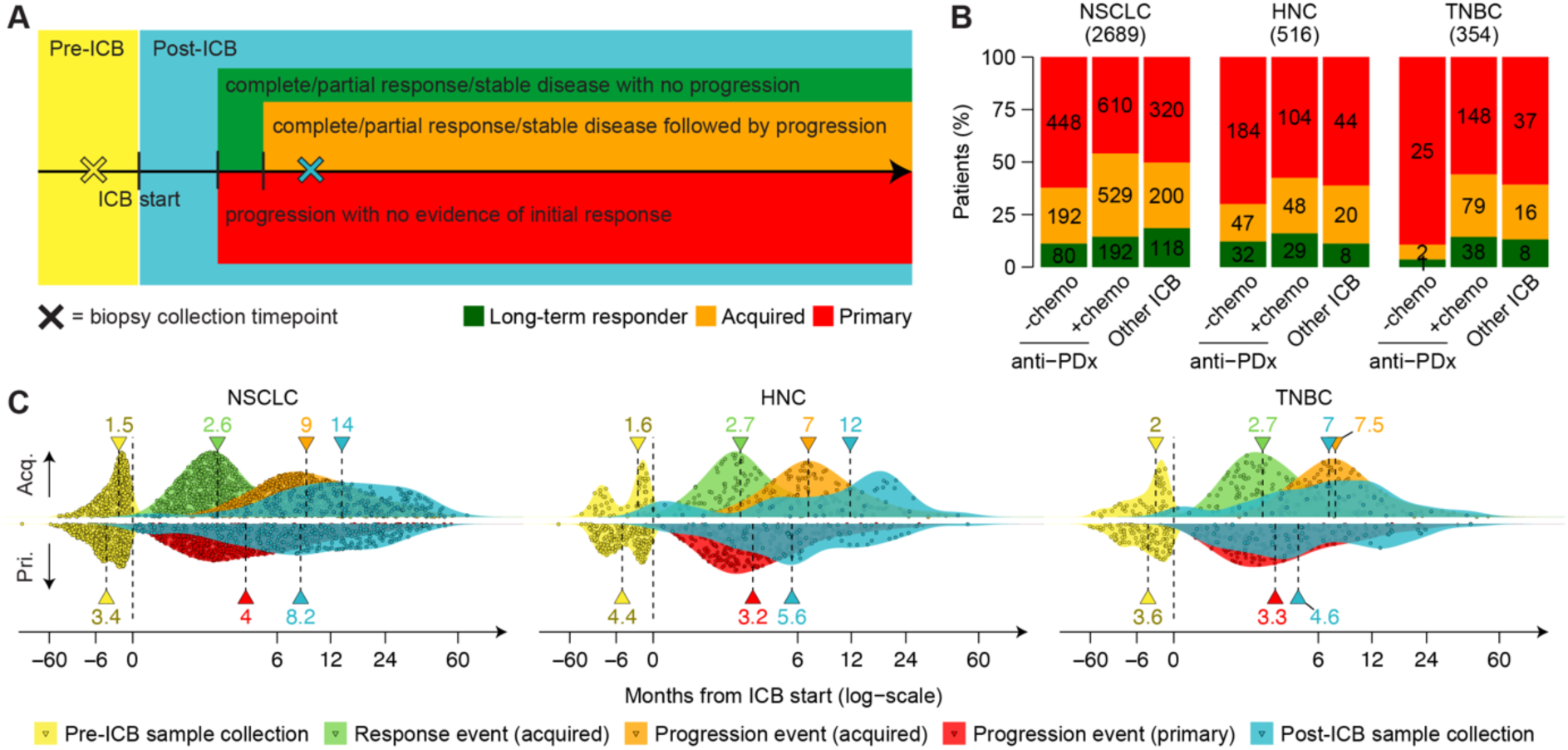
Mapping acquired and primary ICB resistance in real-world data. **A.** Schematic patient timeline showing different timepoints of the treatment journey, including tumour biopsy collection and outcome events. Based on clinical follow-up and outcomes, patients were categorised as long-term responders, acquired or primary resistant to ICB. **B.** Proportion of rwICB long-term responders, acquired, and primary resistant patients across ICB regimens and cancer types. Total number of patients (in brackets) and number of patients in each group are reported. ‘anti-PDx – chemo’ designates anti-PD(L)1 administered as monotherapy, ‘anti-PDx + chemo’ designates anti-PD(L)1 administered concomitantly with chemotherapy, ‘Other ICB’ includes anti-PD1 combined with anti-CTLA-4, anti-PD(L)1 combined with other non-ICB drugs, or any anti-PD(L)1 part of a sequential drug combination. **C.** Collapsed timelines of rwICB acquired (top) and primary (bottom) resistant patients showing distribution of pre/post-ICB sample collection, response events (acquired), and progression events (acquired and primary) relative to ICB start in each cancer type. Each point represents a patient (response/progression events) or a sample (pre/post-ICB sample collection). Dashed lines indicate median months from ICB start for that timepoint. ICB = immune checkpoint blockade, NSCLC = non-small cell lung cancer, HNC = head and neck squamous cell cancer, TNBC = triple-negative breast cancer, anti-PDx = anti-PD(L)1, chemo = chemotherapy.

**Supplementary Figure 1.**
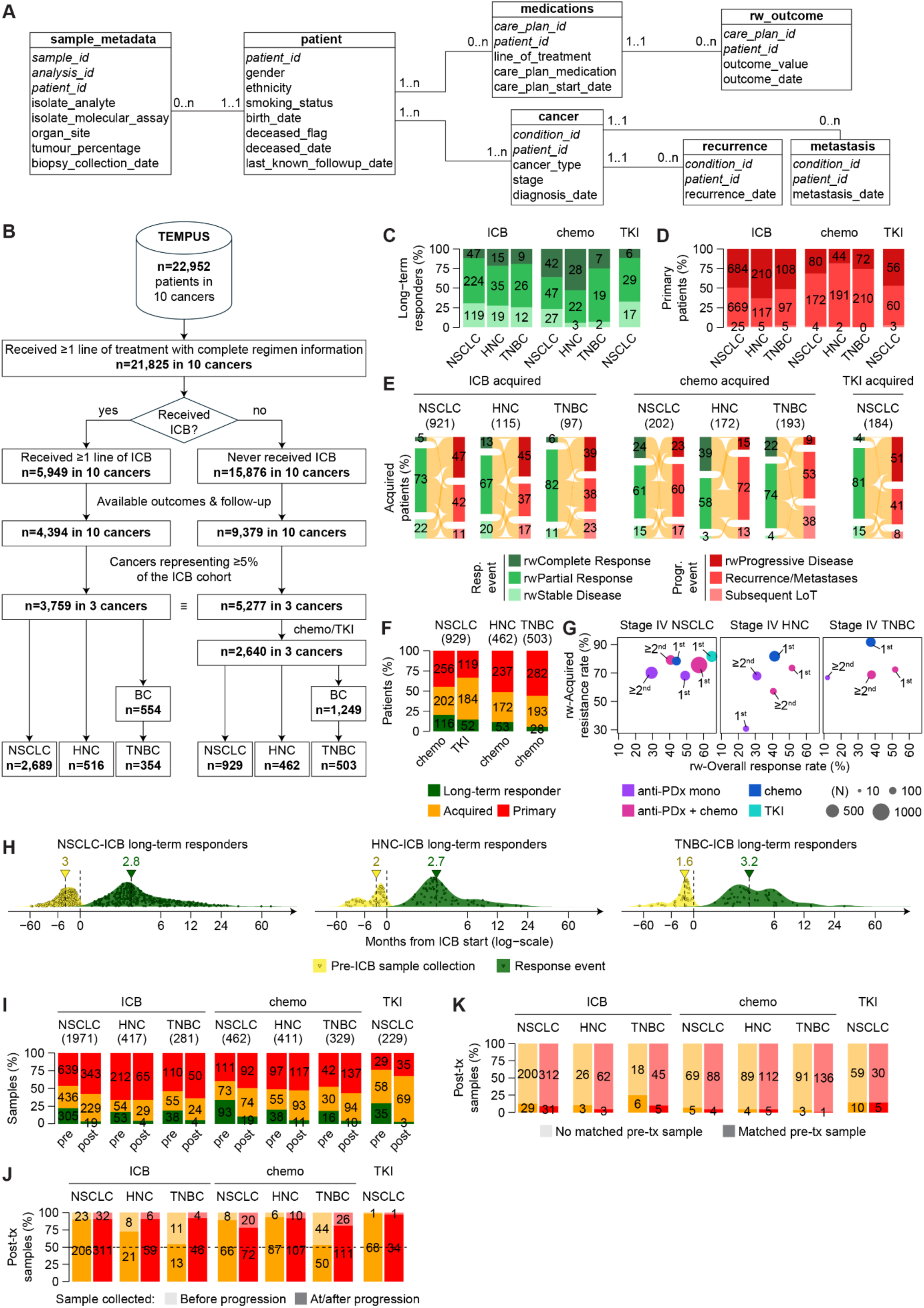
Building cancer treatment datasets from real-world electronic health records. **A.** Entity Relationship Diagram of a simplified version of the multi-modal Tempus AI database. Cardinalities are shown between each pair of linked entities. Primary/secondary keys of each entity are italicised. **B.** Flowchart showing the starting patient total present in the multi-modal Tempus AI database and the resulting numbers at each filtering step throughout the cohort building process. For breast cancer, we focused on the triple-negative subtype where ICB is indicated and in order to exclude subtype-related biases in subsequent analyses. The non-ICB cohort (right hand side) was limited to the corresponding cancer types available in the ICB cohort to serve as a control cohort. **C.** Proportion of long-term responders with rw-Complete/Partial response or durable Stable Disease across treatment modalities and cancer types. **D.** Proportion of primary resistant patients with rw-Progressive Disease, tumour recurrence/metastasis, or a subsequent line of treatment across treatment modalities and cancer types. **E.** Sankey diagrams of initial response (rw-Complete/Partial response or durable Stable Disease) and subsequent progression (rw-Progressive Disease, recurrence/metastasis, or presence of a subsequent line of treatment) for acquired resistant patients across treatment modalities and cancer types. Total number of acquired resistant patients is reported in brackets. Proportion of acquired resistant patients is reported for each type of outcome. **F.** Proportion of long-term responders, acquired or primary resistant patients as annotated in the non-ICB cohort across the three cancer types. Total number of patients is reported in brackets. **G.** Scatter plot of acquired resistance rate vs. overall response rate by treatment modality and line of administration (1^st^ or ≥2^nd^) in stage IV NSCLC, HNC, and TNBC. Circle size is proportional to the total number of patients (long-term responders + acquired + primary resistant patients) in that group. Only treatment groups with N≥10 patients are shown. See Supplementary Table 1 for comprehensive statistics across all (non-)ICB modalities. **H.** Long-term ICB responder patient timeline showing pre-ICB sample collection and response events relative to ICB start in each cancer type. Each point represents a patient (response event) or a sample (pre-ICB sample collection). Dashed lines indicate median months from ICB start for that timepoint. **I.** Proportion of overall DNA/RNA-seq tumour samples available pre/post-treatment in each response group across treatment modalities and cancer types. Total number of samples is reported in brackets. **J.** Proportion of post-treatment tumour samples collected before (light colour shade) or at/after (dark colour shade) progression for acquired and primary resistant patients across treatment modalities and cancer types. **K.** Proportion of post-treatment tumour samples with (dark colour shade) or without (light colour shade) a matched pre-treatment sample for acquired and primary resistant patients across treatment modalities and cancer types. Number of patients/samples is reported for each group in **C**, **D**, **F**, **I**, **J**, **K**. rw = real-world, ICB = immune checkpoint blockade, Resp. = responder, Progr. = progression, NSCLC = non-small cell lung cancer, HNC = head and neck squamous cell cancer, BC = breast cancer, TNBC = triple-negative breast cancer, LoT = line of treatment, anti-PDx = anti-PD(L)1, chemo = chemotherapy, mono = monotherapy, TKI = tyrosine kinase inhibitors, tx = treatment.

Overall, rwICB reflected expectations of outcome to ICB, with <15% of patients showing long-term benefit from treatment, >50% experiencing primary resistance, and ∼22-34% experiencing acquired resistance, varying across cancer types and treatment settings (Figure 1B, Supplementary Figure 1F). Expectedly, addition of chemotherapy to the ICB regimen improved the real-world overall response rate (i.e. the proportion of patients with long-term response or initial response followed by acquired resistance) across all cancer types^2,4^ (Figure 1B). However, the proportion of patients experiencing acquired resistance was consistently higher for those treated with chemoimmunotherapy (27-40%) as compared to those treated with anti-PD(L)1 monotherapy (7-27%, Figure 1B, Supplementary Figure 1G, Supplementary Table 1).

In rwICB, initial response to ICB occurred mostly within six months of treatment start for both long-term responders (>76%) and acquired resistant patients (>87%), while progression was reported within a year of treatment start for most acquired resistant patients (66% in NSCLC, >80% in HNC and TNBC) and within six months of treatment start for most primary resistant patients (67% in NSCLC, 79% in HNC, and 72% in TNBC, Figure 1C, Supplementary Figure 1H, Supplementary Table 1). In addition to annotating patient outcomes, we used biopsy collection dates from the database to annotate pre– and post-treatment tumour DNA/RNA sample availability across all cohorts (Supplementary Figure 1A, Supplementary Figure 1I). Nearly all post-treatment samples came from acquired or primary resistant patients, reflecting absence of tumour tissue in long-term responders due to effective tumour clearance following treatment. Most post-ICB resistant tumours were collected at or after progression for both acquired and primary resistant patients, while the majority of pre-ICB samples were collected up to six months before treatment start (Figure 1C, Supplementary Figure 1H, Supplementary Figure 1J). Importantly, pre– and post-treatment samples in these cohorts were typically unmatched (i.e. they come from different patients, Supplementary Figure 1K).

### Clinical features of acquired and primary resistance to ICB

Using the comprehensive clinicogenomic data available in rwICB, we sought to compare the baseline patient profiles of our three defined response groups across treatment modalities. First, we computed and compared the real-world overall survival (rwOS) across response groups. To mitigate survival overestimation inherent to real-world clinicogenomic databases^21^, we performed risk set adjustment in all survival analyses using tumour sequencing date to account for delayed entries (Methods). We confirmed that rwOS following treatment was longest for long-term responders, followed by acquired then primary resistant patients, across all cancer types and regardless of stage or treatment modality (Supplementary Figures 2A-B, Supplementary Table 2). We then compared a set of established ICB biomarkers^4^ along with other clinical variables across patient groups (Figure 2A, Supplementary Table 3). In NSCLC, higher TMB, PD-L1 protein expression, and inferred immune infiltration associated with long-term benefit to ICB, while only higher PD-L1 expression and immune infiltration associated also with acquired vs. primary ICB resistance. Interestingly, chemotherapy NSCLC responders also showed higher TMB and PD-L1 expression at baseline (Supplementary Table 3), suggesting these biomarkers are not ICB-specific. In HCN and TNBC, there were no clear differences in TMB, PD-L1 expression, or immune infiltration between acquired vs. primary ICB resistance. This suggests that currently used response biomarkers cannot generally predict acquired resistance to ICB.

**Figure 2.**
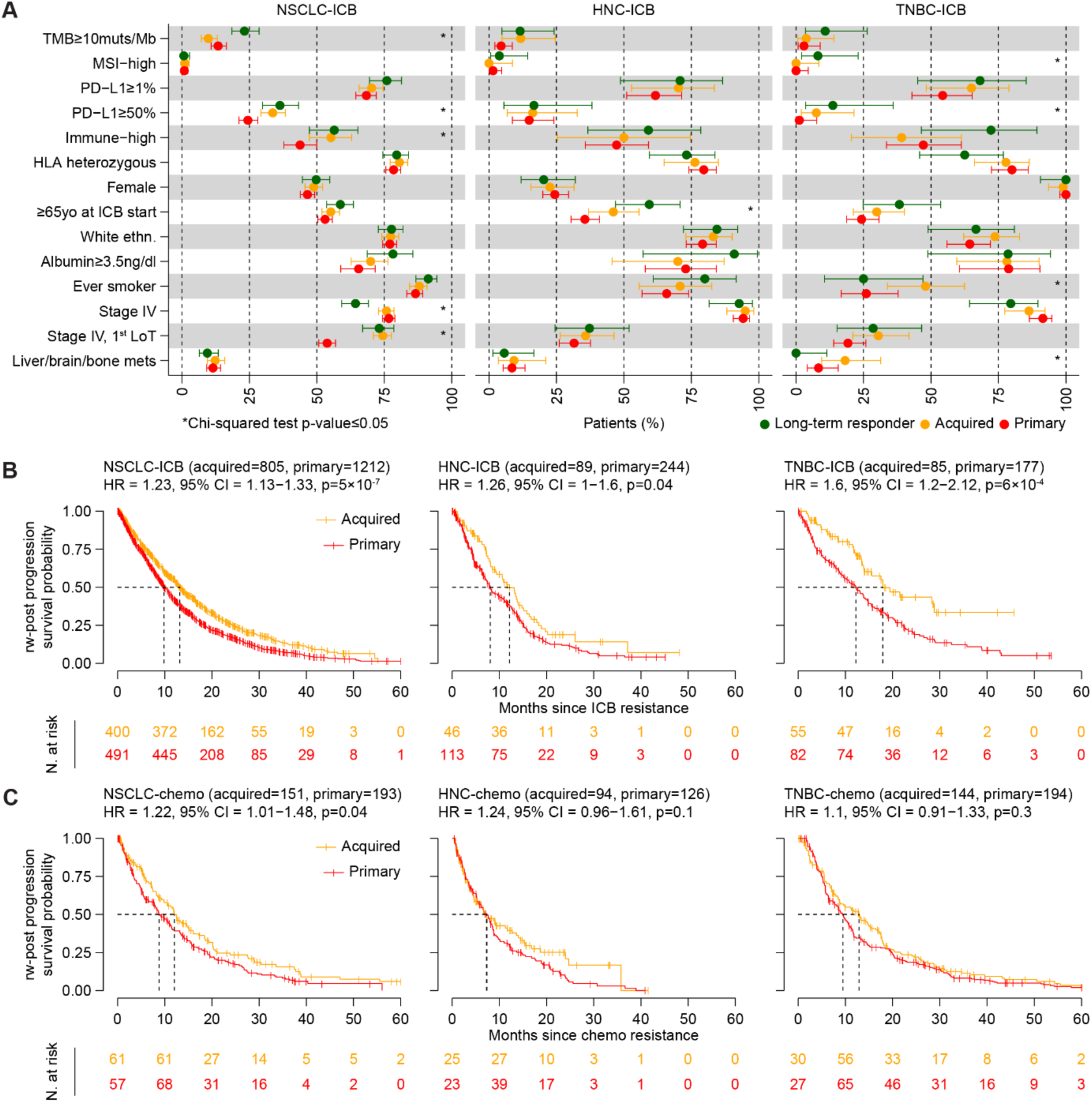
ICB prolongs survival post-acquired resistance. **A.** Proportions of ICB long-term responders, acquired and primary resistant patients across a range of baseline (pre-treatment) clinical features in each cancer type. Error bars represent the 95% confidence interval of each proportion as computed using the one-sample proportions test with ‘prop.test’ R function. PD-L1 protein expression was assessed via immunohistochemistry and the readout was calculated using the tumour proportion score (TPS) approach. Immune-high designates an Immune Score^23^ above the median within each cancer type as calculated using single sample gene set enrichment analysis^24^ on RNA-seq data (Methods, only tumours biopsied from the corresponding primary organ site were used. In HNC only tumours with purity ≥50% were used due to baseline differences (Supplementary Figure 2C)). HLA class I haplotypes were derived from DNA-seq data and ‘HLA heterozygous’ refers to heterozygosity at all three loci (HLA-A/B/C). Presence of liver/brain/bone mets at baseline was determined using pre-ICB tumour samples from rwICB (Supplementary Figure 1I). Asterisks indicate statistical significance (p-value≤0.05) comparing the three patient groups with the Chi-squared test. Additional features, breakdown across (non-)ICB treatment modalities, and statistical comparisons of acquired vs. primary resistance are reported in Supplementary Table 3. Risk set-adjusted Kaplan-Meier curves of the real-world five-year post-progression survival probability for acquired and primary ICB-**(B)** or chemotherapy-**(C)** resistant patients in each cancer type. The total number of patients used in the analysis is reported in brackets for each group. Hazard ratios, corresponding 95% confidence intervals, and likelihood ratio test p-values comparing acquired vs. primary resistant patients are reported. Number of patients at risk at ten-month intervals is reported for each group. Only patients with complete last known follow-up and censoring information are included in the analysis. Note that the total number of patients is different from the number of patients at risk at T=0 because of delayed entry due to risk set adjustment (Methods). Analyses stratified by (non-)ICB treatment modalities and disease stage are reported in Supplementary Table 2. NSCLC = non-small cell lung cancer, HNC = head and neck squamous cell cancer, TNBC = triple-negative breast cancer, TMB = tumour mutational burden, muts/Mb = mutations per mega bases, MSI = microsatellite instability, HLA = human leukocyte antigen, ICB = immune checkpoint blockade, ethn. = ethnicity, LoT = line of treatment, mets = metastases, HR = hazard ratio, CI = confidence interval, rw = real-world, chemo = chemotherapy, n.s. = not significant.

**Supplementary Figure 2.**
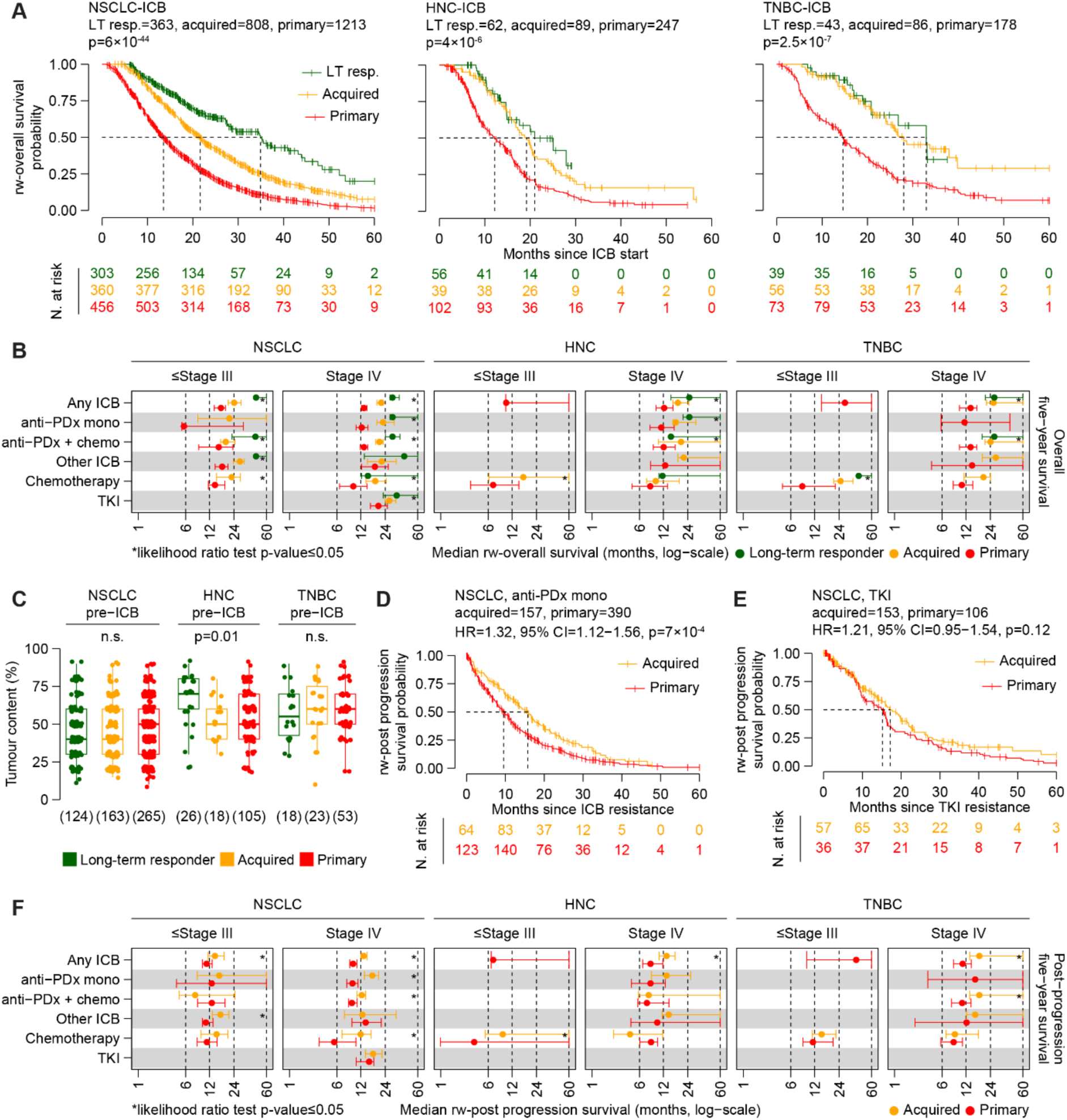
Prognostic analysis of patient groups across treatment settings. **A.** Risk set-adjusted Kaplan-Meier curves of the real-world five-year overall survival probability for long-term responders, acquired and primary resistant patients in each cancer type of the ICB cohort. The total number of patients, number of patients at risk at ten-month intervals, and likelihood ratio test p-values comparing the three patient groups are reported. Note that the total number of patients is different from the number of patients at risk at T=0 because of delayed entry due to risk set adjustment (Methods). **B.** Risk set-adjusted median real-world five-year overall survival for long-term responders, acquired and primary resistant patients stratified by treatment modality and disease stage in each cancer type. Error bars represent the 95% confidence interval of the median. Asterisks indicate statistical significance (p-value ≤0.05) comparing the three patient groups with the likelihood ratio test. Number of patients in each group, medians, and p-values are reported in Supplementary Table 2. **C.** Distribution of the tumour content in pre-ICB tumour RNA samples across patient groups in each cancer type. Only primary site samples are considered (lung tissue for NSCLC, head and neck tissue for HNC, and breast tissue for TNBC). Number of samples in each group is reported in brackets. Tumour content distributions were compared with the Kruskal–Wallis test across patient groups within each cancer type. Risk set-adjusted Kaplan-Meier curves of the real-world five-year post-progression survival probability for acquired and primary anti-PDx mono-**(D)** or TKI-**(E)** resistant NSCLC patients. The total number of patients used in the analysis is reported for each group. Hazard ratios, corresponding 95% confidence intervals, and likelihood ratio test p-values comparing acquired vs. primary resistant groups are reported. Number of patients at risk at ten-month intervals is reported for each group. Note that the total number of patients is different from the number of patients at risk at T=0 because of delayed entry due to risk set adjustment (Methods). Analyses stratified by (non-)ICB treatment modalities and disease stage are reported in Supplementary Table 2. **F.** Risk set adjusted median real-world five-year post-progression survival for acquired and primary resistant patients stratified by treatment modality and disease stage in each cancer type. Error bars represent the 95% confidence interval of the median. Asterisks indicate statistical significance (p-value≤0.05) comparing acquired vs. primary resistant patients with the likelihood ratio test. Number of patients in each group, medians, and p-values are reported in Supplementary Table 2. For **A**, **B**, **D**, **E** and **F**, only patients with complete last known follow-up and censoring information were included in the analysis. For **B** and **F**, only groups with N≥10 patients and whose median survival is reached (i.e. ≠ N/A) are shown. See Supplementary Table 2 for all survival analysis results. NSCLC = non-small cell lung cancer, HNC = head and neck squamous cell cancer, TNBC = triple-negative breast cancer, ICB = immune checkpoint blockade, LT resp. = long-term responders, anti-PDx = anti-PD(L)1, mono = monotherapy, chemo = chemotherapy, TKI = tyrosine kinase inhibitors, rw = real-world.

Patient-level characteristics were generally similar across response groups, including germline zygosity of the Human Leukocyte Antigen (consistent with^22^), patient gender, or ethnicity (Figure 2A, Supplementary Table 3). Older age associated with durable ICB benefit, particularly in HNC, while smoking history and presence of distant metastases pre-ICB associated specifically with acquired resistance in TNBC. Despite ICB long-term responders tending to present with earlier stage disease, there was no difference in disease stage between acquired and primary resistant patients. However, among stage IV patients, those who experienced acquired resistance were more likely to have received ICB as their first ever treatment compared to primary resistant patients, especially in NSCLC as previously reported^8^.

Focusing only on acquired or primary resistant patients, we compared them prognostically after they progressed on ICB by calculating their risk set-adjusted real-world post-progression survival (rwPPS, i.e. overall survival calculated from the onset of progression). Despite equally advanced stage, acquired ICB resistant patients showed significantly longer rwPPS than primary resistant patients across the three cancer types (Figure 2B). In NSCLC, this observation was regardless of line of ICB administration (Supplementary Table 2) and was even more striking in patients treated with anti-PD-(L)1 monotherapy (Supplementary Figure 2D), which aligns with a previous report in the same patient setting^8^. Interestingly, this prognostic difference was not observed in chemotherapy– or TKI-treated patients (Figure 2C, Supplementary Figure 2E), even when stratifying by stage (Supplementary Figure 2F), as acquired generally survived as long as their primary counterpart post-progression. These results indicate a universal and ICB-specific prognostic advantage procured by an initial response that, although less effective, may still sustain patient survival after acquiring resistance. This therefore suggests ICB-specific differences at the molecular level between acquired and primary resistant patients.

### The acquired resistant immune TME is consistently inflamed post-ICB

Given the consistently prolonged survival in acquired vs. primary ICB resistant patients following progression, we hypothesised that differences at the molecular level underpinned these distinct clinical phenotypes. We leveraged the multi-modal samples available in rwICB and sought to first compare the transcriptional profiles of acquired vs. primary resistant tumours post-ICB. We used RNA-seq data of tumours biopsied from the corresponding primary organ site of each cancer type (lung tissue for NSCLC, head/neck tissue for HNC, and breast tissue for TNBC) to prevent introducing variability due to distinct metastatic sites (Supplementary Figure 3A). Since RNA-seq was performed with two different assays (Supplementary Figure 3B), we accounted for the assay version as a co-variate in the differential expression analysis (DEA) for all cancer types, while also accounting for histology in NSCLC. To estimate the immune composition of the TME and the level of oncogenic signalling across tumour groups, we ran gene set enrichment analysis (GSEA) on the DEA-ranked list of genes using 50 Hallmark gene sets from MSigDB^25^ and 19 cancer-specific immune signatures from ConsensusTME^23^ (Methods, GSEA results reported in Supplementary Table 4).

Post-ICB, acquired resistant NSCLCs showed a significantly immune-inflamed TME as compared to primary resistant tumours (Figure 3A). This immune-inflamed profile was characterised by inferred infiltration of different lymphocytes and myeloid cells, as well as active IFNγ signalling and inflammatory responses. These immune differences were more prevalent in anti-PD(L)1 monotherapy-treated NSCLC or in the ≥2^nd^ line advanced-stage setting (Supplementary Table 4). Remarkably, this immunophenotype was consistent across all cancer types, with acquired resistant HNCs and TNBCs also exhibiting increased immune infiltration and IFNγ signalling as compared to primary resistant tumours post-ICB (Figure 3B). We checked that these results were not confounded by differences in TMB (Figure 3C), PD-L1 protein expression (Figure 3D), or tumour purity (Supplementary Figure 3C). We also checked that timing of tumour biopsy with respect to both ICB start and ICB progression did not correlate with overall immune infiltration in either acquired or primary resistant tumours (Supplementary Figures 3D-E). Since squamous and non-squamous NSCLCs showed markedly distinct transcriptional profiles (Supplementary Figure 3F), we stratified DEA and GSEA by histology in NSCLC to understand if immune profiles post-ICB resistance were histology-dependent or widespread. Again, acquired resistant NSCLCs showed an immune-inflamed profile vs. primary resistant tumours post-ICB, regardless of histology (Supplementary Figure 3G). Interestingly, the main histology-related discrepancies pertained to oncogenic signalling pathways, including differential enrichment of Myc, KRAS, or epithelial mesenchymal transition (EMT) pathways depending on histology (Supplementary Figure 3G).

**Figure 3.**
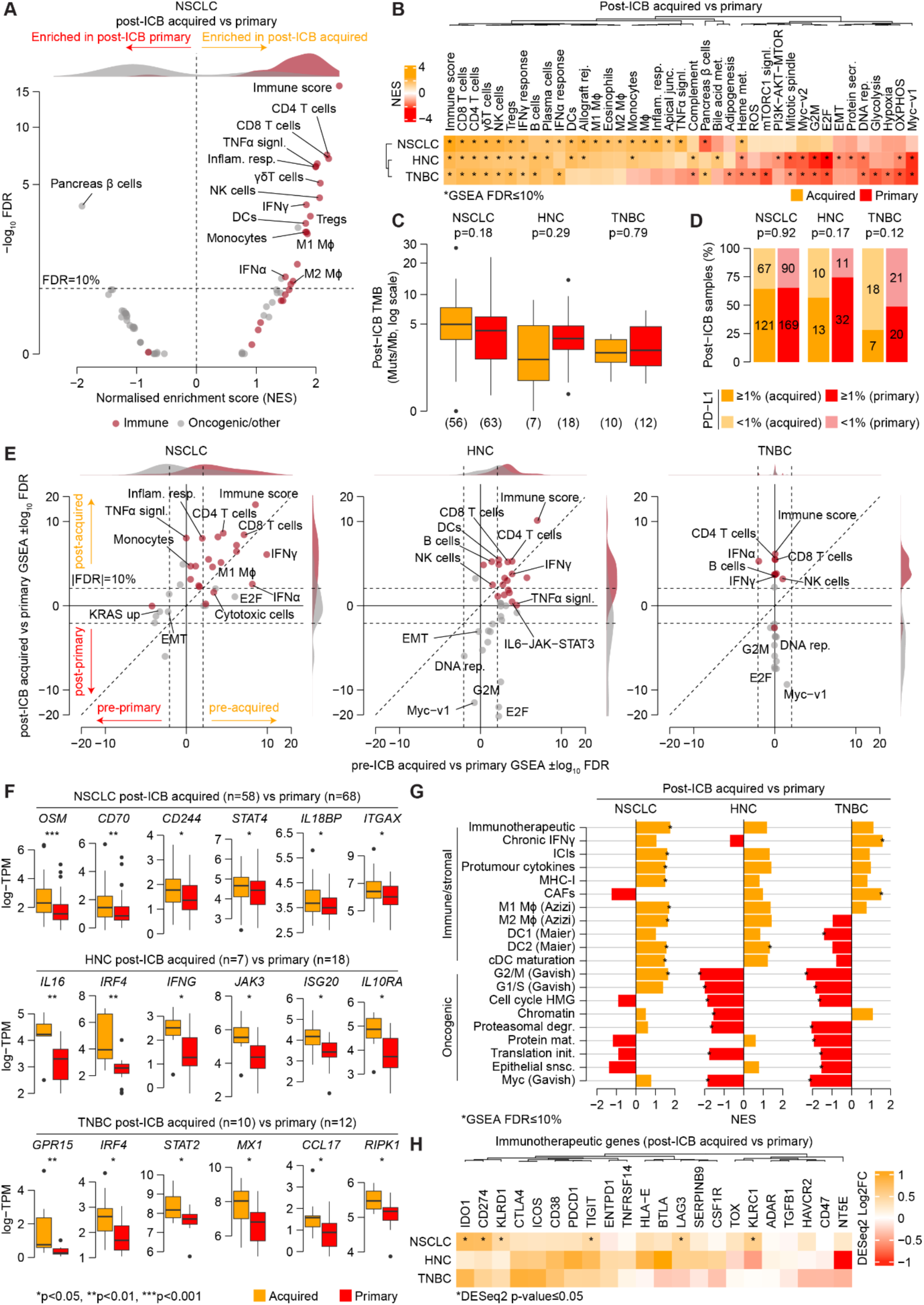
ICB maintains or induces an immune-inflamed TME post-acquired resistance. **A.** Volcano plot of differentially enriched gene sets in post-ICB acquired vs. primary resistant NSCLCs. Gene sets are coloured by functional category. NES density curves for immune and non-immune gene sets are shown at the top. Horizontal dashed line indicates significance at FDR=10%. **B.** GSEA NES heatmap of differentially enriched gene sets in post-ICB acquired vs. primary resistance across the three cancer types. Rows and columns are clustered hierarchically. **C.** TMB distribution in post-ICB acquired vs. primary resistant tumour samples used in RNA-seq analysis. Distributions are compared within each cancer type using Wilcoxon rank sum test. Number of samples in each group is reported in brackets. In NSCLC, two acquired resistant samples and five primary resistant samples had no corresponding TMB as assessed using xTv2, xTv3, or xTv4 DNA-seq assays (Methods). **D.** Proportion of post-ICB acquired vs. primary resistant tumours with PD-L1 protein expression ≥1%. PD-L1 protein expression was assessed via immunohistochemistry and the readout was calculated using the tumour proportion score (TPS) approach. Proportions are compared within each cancer type using Fisher’s exact test. Number of samples is reported for each group. All post-ICB samples with available PD-L1 data were used. **E.** Scatter plots of differentially enriched gene sets in pre– or post-ICB acquired vs. primary resistance across the three cancer types. For each gene set, the NES-signed GSEA log_10_ FDR for the pre-ICB and post-ICB comparison is reported on the x and y axis respectively, with positive values indicating enrichment in pre/post acquired, and negative values indicating enrichment in pre/post primary. Gene sets are coloured by functional category. ±Log_10_ FDR density curves for immune and non-immune gene sets are shown at the top for pre-ICB comparisons and on the right for post-ICB comparisons. Diagonal dashed lines indicate the identity line. Horizontal and vertical dashed lines indicate significance at FDR=10%. **F.** Log-transcript per million (TPM) distribution of representative immune-related genes in post-ICB acquired vs. primary resistant tumours across the three cancer types. Number of samples in each group is reported in brackets. Distributions are compared for each gene within each cancer type using Wilcoxon rank sum test. **G.** GSEA NES barplots of differentially enriched literature-compiled gene sets in post-ICB acquired vs. primary resistance across the three cancer types. Positive NES values indicate enrichment in post-ICB acquired, negative NES values indicate enrichment in post-ICB primary. Gene sets are grouped by functional category. **H.** Post-ICB acquired vs. primary DESeq2 log2-fold change (FC) heatmap of genes from the immunotherapeutic gene set^8^ across the three cancer types. Columns are clustered hierarchically. FDR was computed using Benjamini-Hochberg approach in all analyses. For **B**, **E**, and **G**, only gene sets differentially enriched in at least one comparison in any cancer type are shown. See Supplementary Table 4 for gene set abbreviations and all GSEA results. NSCLC = non-small cell lung cancer, HNC = head and neck squamous cell cancer, TNBC = triple-negative breast cancer, ICB = immune checkpoint blockade, GSEA = gene set enrichment analysis, NES = normalised enrichment score, FDR = false discovery rate, TMB = tumour mutational burden, muts/Mb = mutations per mega bases, TPM = transcripts per million, FC = fold change.

**Supplementary Figure 3.**
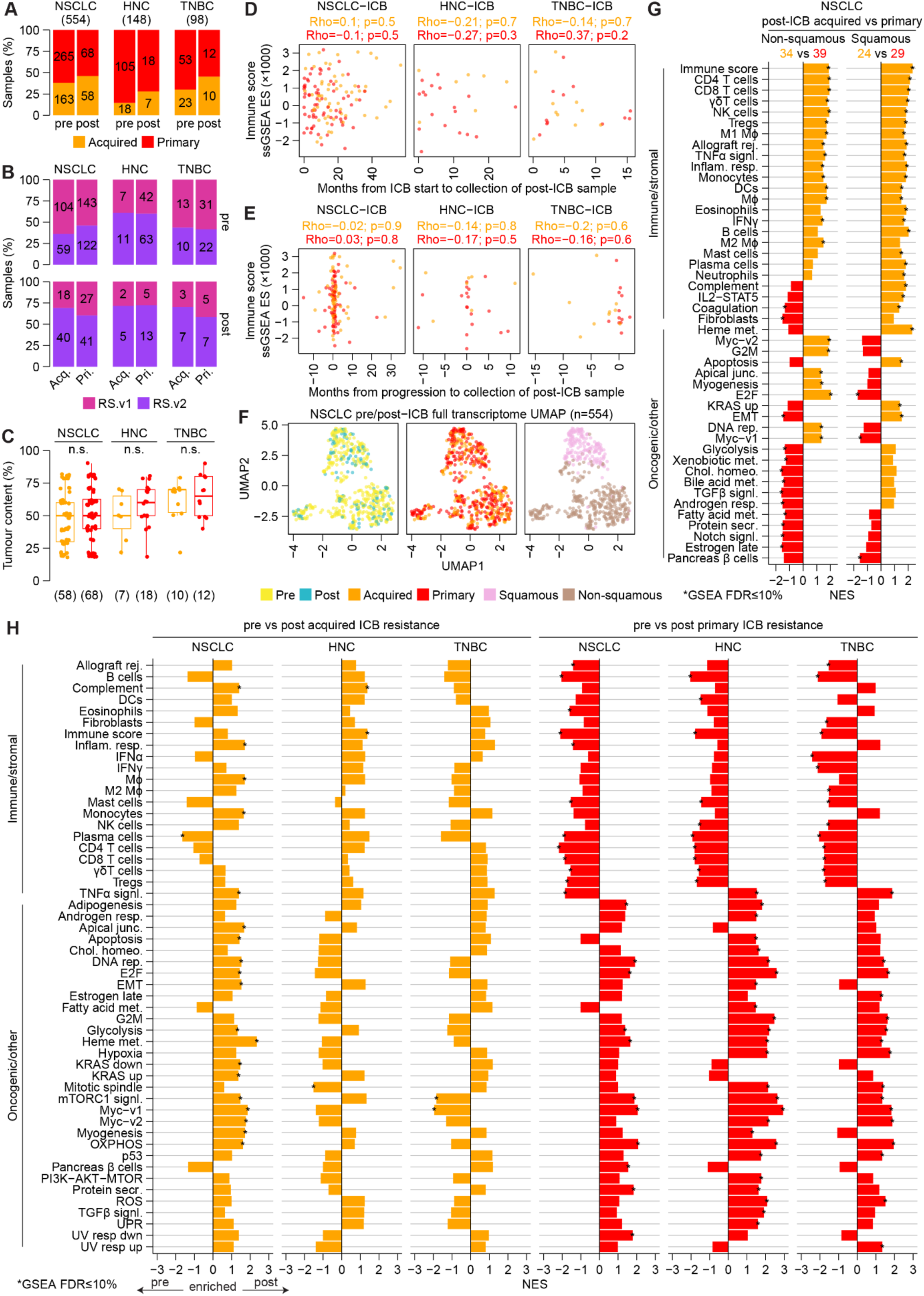
Transcriptomic profiling of pre/post-ICB acquired vs. primary resistant tumours. **A.** Proportion of pre/post-ICB RNA-seq tumour samples biopsied from the primary organ site of each cancer type in acquired and primary resistant patients. **B.** Proportion of pre/post-ICB samples from acquired/primary resistant patients whose tumour was sequenced for RNA with the RS.v1 or RS.v2 Tempus AI assays across the three cancer types. **C.** Distribution of tumour content (purity) in post-ICB acquired vs. primary resistant samples used in RNA-seq analysis. Distributions are compared within each cancer type using Wilcoxon rank sum test. Number of samples in each group is reported in brackets. Correlation analysis of overall immune infiltration vs. timing of post-ICB RNA-seq sample collection with respect to **(D)** ICB start or **(E)** ICB progression for acquired and primary resistant patients. Spearman correlation score (Rho) and associated p-value are reported for both acquired and primary resistant samples in each cancer type. Overall immune infiltration was estimated via ssGSEA of the Immune Score signature from ConsensusTME^23^. Number of post-ICB samples in each group corresponds to that reported in panel **A**. **F.** UMAP dimensionality reduction on the full transcriptome of pre/post-ICB NSCLC RNA-seq samples from acquired and primary resistant patients. Total number of samples is reported in brackets. Samples are coloured by pre/post-ICB timepoint, acquired/primary resistance, or squamous/non-squamous histology. Only samples biopsied from the primary organ site (lung tissue) were used. **G.** GSEA NES barplots of differentially enriched gene sets in post-ICB acquired vs. primary resistant non-squamous and squamous NSCLC, respectively. Positive NES values indicate enrichment in post acquired, negative NES values indicate enrichment in post primary. Number of samples in each group is reported for both comparisons. **H.** GSEA NES barplots of differentially enriched gene sets in pre-vs. post-ICB acquired or primary resistance across the three cancer types. Positive NES values indicate enrichment in post acquired or post primary, negative NES values indicate enrichment in pre-ICB acquired or pre-ICB primary. FDR was computed using Benjamini-Hochberg approach in all analyses. In **A** and **B**, Number of samples is reported for each group. For **G** and **H**, gene sets are grouped by functional category and only those differentially enriched in at least one comparison in any cancer (sub)type are shown. See Supplementary Table 4 for gene set abbreviations, number of samples in each group, and all GSEA results. NSCLC = non-small cell lung cancer, HNC = head and neck squamous cell cancer, TNBC = triple-negative breast cancer, ICB = immune checkpoint blockade, Acq. = acquired, Pri. = primary, n.s. = not significant, ssGSEA = single sample gene set enrichment analysis, ES = enrichment score, UMAP = Uniform Manifold Approximation and Projection, GSEA = gene set enrichment analysis, NES = normalised enrichment score, FDR = false discovery rate.

To evaluate the post-ICB specificity of the immune-inflamed acquired resistant TME, and since most post-treatment samples do not have a matched pre-treatment sample (Supplementary Figure 1K), we contrasted post-ICB GSEA results with pre-ICB comparisons. While pre-ICB acquired resistant NSCLCs and HNCs were also more immune-inflamed than pre-ICB primary resistant tumours, immune-related differences were exacerbated in, or sometimes exclusive to, the post-ICB setting across all cancer types (Figure 3E). TNBC stands as a typical case, where no transcriptional differences were found between acquired and primary resistant tumours pre-ICB, yet a substantially different TME was observed post-ICB (Figure 3E). Despite coming from different patients, we also compared pre-vs. post-ICB tumours from acquired and primary resistant patients respectively. In line with our previous analysis of paired pre/post-ICB acquired resistant NSCLCs^8^, we found stable or increased immune inflammation in acquired resistant tumours from pre-to post-ICB, including increased inflammatory response and tumour necrosis factor alpha (TNFα) signalling in NSCLC (Supplementary Figure 3H). Contrastingly, primary resistant tumours consistently showed vigorous immune depletion from pre-to post-ICB, further supporting a fundamentally different post-ICB TME in the two resistance phenotypes. Moreover, no immune differences were found in post-chemotherapy acquired vs. primary resistant HNCs or TNBCs, while post-chemotherapy (but not post-TKI) acquired resistant NSCLCs were moderately more inflamed than primary resistant tumours (Supplementary Table 4). Taken together, these results indicate that post-ICB immunophenotypic differences between acquired and primary resistance are not merely a reflection of pre-existing baseline differences, but are rather specifically induced by ICB.

Different genes driving these immune differences were upregulated in post-ICB acquired vs. primary resistant tumours across the three cancer types (Figure 3F, Supplementary Table 4). These include genes encoding components of immune inflammatory processes (*OSM*, *CD70*, *IL16*, *CCL17*), interferon signalling (*STAT2/4*, *IL18BP*, *IRF4*, *IFNG*, *JAK3*, *ISG20*, *MX1*, *RIPK1*), or immune cell markers (*CD244*, *ITGAX*, *IL10RA*, *GPR15*). To further characterise the post-ICB acquired resistant immunophenotype, we ran GSEA on a complementary list of immuno-oncology-focused gene sets we compiled from the literature^8,26–29^. Gene sets describing immunotherapy targets, alternative immune checkpoints, pro-tumour cytokines, and antigen processing and presentation components were all enriched in post-ICB acquired vs. primary resistant NSCLCs, with similar trends in HNC and TNBC (Figures 3G-H). Cancer-related processes, including cell proliferation and Myc signalling, were on the other hand mainly enriched in primary resistant tumours, particularly in HNC and TNBC (Figures 3G).

Altogether, our transcriptional profiling of post-ICB resistant tumours revealed a universally immune-inflamed TME in the acquired resistance setting across the three cancer types studied, suggestive of an ongoing, albeit impaired, ICB-induced immune response.

### ICB-specific selection of mutations in the post-acquired resistance setting

The distinctive immunophenotype of acquired resistant tumours post-ICB prompted us to probe their genetic landscape for presence of ICB-specific cancer alterations that may confer resistance to treatment. To this end, we leveraged panel DNA-seq data covering 592 cancer-related genes and used *dndscv*^30^ to quantify the level of selective pressure acting on gene mutations from pre-to post-treatment. Briefly, *dndscv* identifies genes undergoing positive selection by quantifying the ratio of non-synonymous to synonymous mutations (dN/dS) using local (same gene, cross-samples) and global (cross-genes) information to estimate the background mutation rate. This approach presents many advantages, including sensitivity to low-frequency events and robustness against potential confounding factors in cancer genomics data, such as mutational signature biases and variability in tumour purity and/or TMB^30^.

We used cross-organ site samples to run *dndscv* in each of pre-/post-treatment acquired/primary resistant tumour groups separately within each cancer type and for each of ICB, chemotherapy, and TKI treatment modalities (Figure 4A, Supplementary Figure 4A, Supplementary Table 5). To exclude biases, we checked sample distribution by sequencing panel, NSCLC histology, organ site, and availability of matched tumour/normal sample (germline control) for DNA-seq (Supplementary Figures 4B-E). We also checked that the global dN/dS estimates in the mutation datasets were ≥1 (Supplementary Figure 4F), as expected^30^.

**Figure 4.**
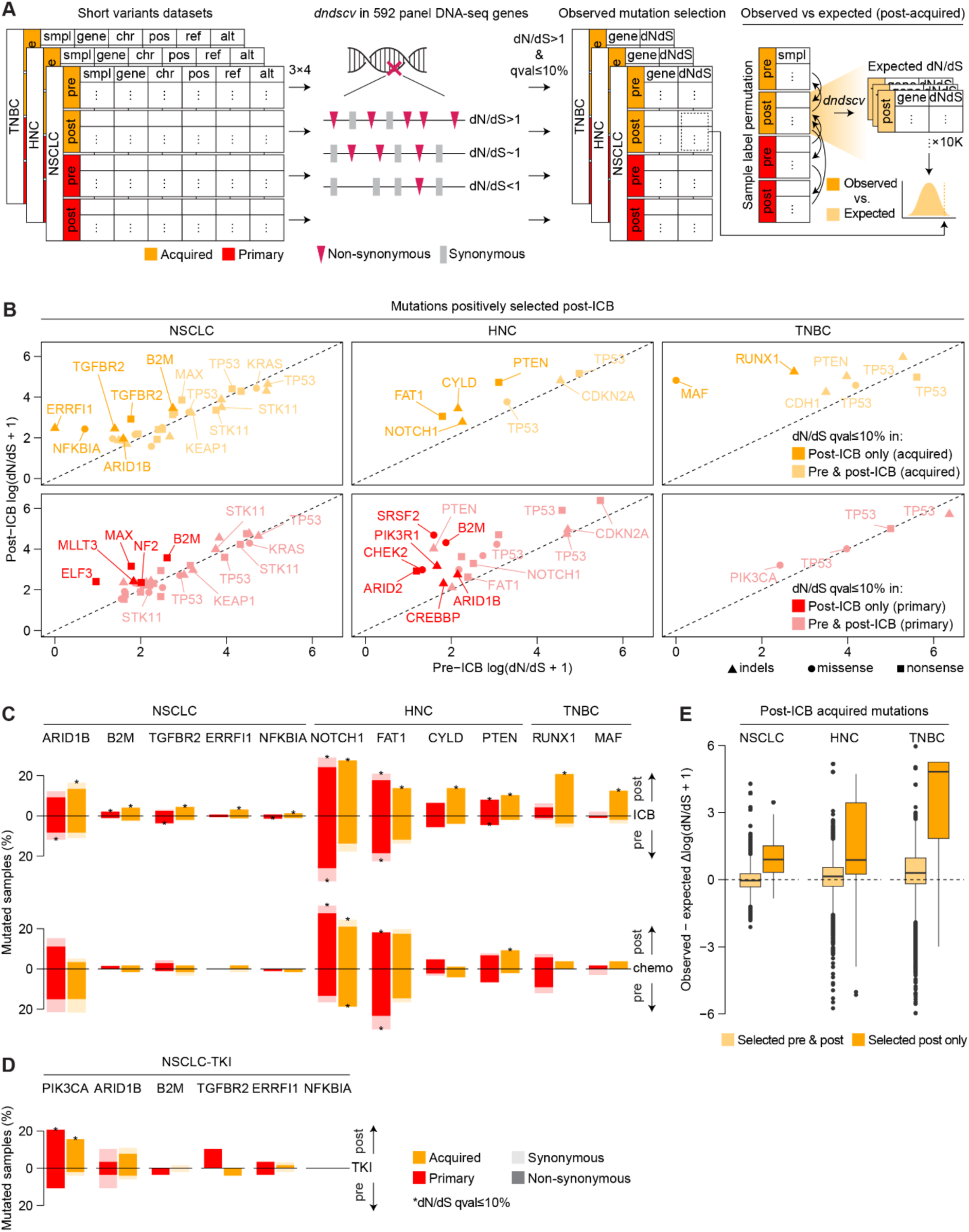
ICB selects for mutations specific to the acquired resistance setting. **A.** Schematic of *dndscv* analysis. For each cancer type and treatment modality (ICB, chemo, TKI), *dndscv* was run separately on short variant data of pre-/post-treatment acquired/primary resistant tumours to identify mutations under significant positive selection (dN/dS>1 and qval≤10%). Observed dN/dS estimates of positively selected mutations in post-ICB acquired resistance were further compared against an expected distribution of dN/dS estimates generated by randomly shuffling pre/post acquired/primary tumour group labels within each cancer type of the ICB cohort. **B.** Scatter plot of pre-ICB (x-axis) and post-ICB (y-axis) dN/dS estimates of all positively selected post-ICB acquired (upper panels) and primary (lower panels) resistance mutations across the three cancer types. Darker colour shade indicates mutations identified post-ICB only, while lighter shade indicates mutations identified also pre-ICB (i.e. both pre– and post-ICB). Dot shape indicates mutation type (missense, nonsense, indels). Dashed lines represent the identity line (i.e. pre-ICB dN/dS = post-ICB dN/dS). Stacked barplots showing the recurrence of positively selected post-ICB acquired resistance mutations in acquired and primary resistant tumours pre– and post-ICB/chemo **(C)** or TKI **(D)** across the three cancer types. Recurrence of both non-synonymous (darker colour shade) and synonymous (lighter colour shade) mutations is reported. Asterisks indicate mutations positively selected in that tumour group. **E.** Distribution of the difference between the observed and expected dN/dS estimates (ΔdN/dS) for post-ICB acquired resistance mutations positively selected both pre– and post-ICB (light orange) or post-ICB only (dark orange) in each cancer type of the ICB cohort. Each dot represents a gene mutation and each gene mutation appears 10,000 times, each time corresponding to one iteration of the random shuffling process performed to get an expected dN/dS estimate (Methods). ΔdN/dS values >0 indicate higher-than-expected dN/dS estimates. For **C** and **D**, the total number of samples in each tumour group (i.e. denominator) is reported in Supplementary Figure 4A. ‘Non-synonymous’ refers to any non-synonymous substitution or indel. For **D**, *PIK3CA* is shown as a TKI positive control alongside post-ICB acquired mutations. NSCLC = non-small cell lung cancer, HNC = head and neck squamous cell cancer, TNBC = triple-negative breast cancer, smpl = sample, chr = chromosome, pos = position, ref = reference allele, alt = alternative allele, ICB = immune checkpoint blockade, chemo = chemotherapy, TKI = tyrosine kinase inhibitors.

**Supplementary Figure 4.**
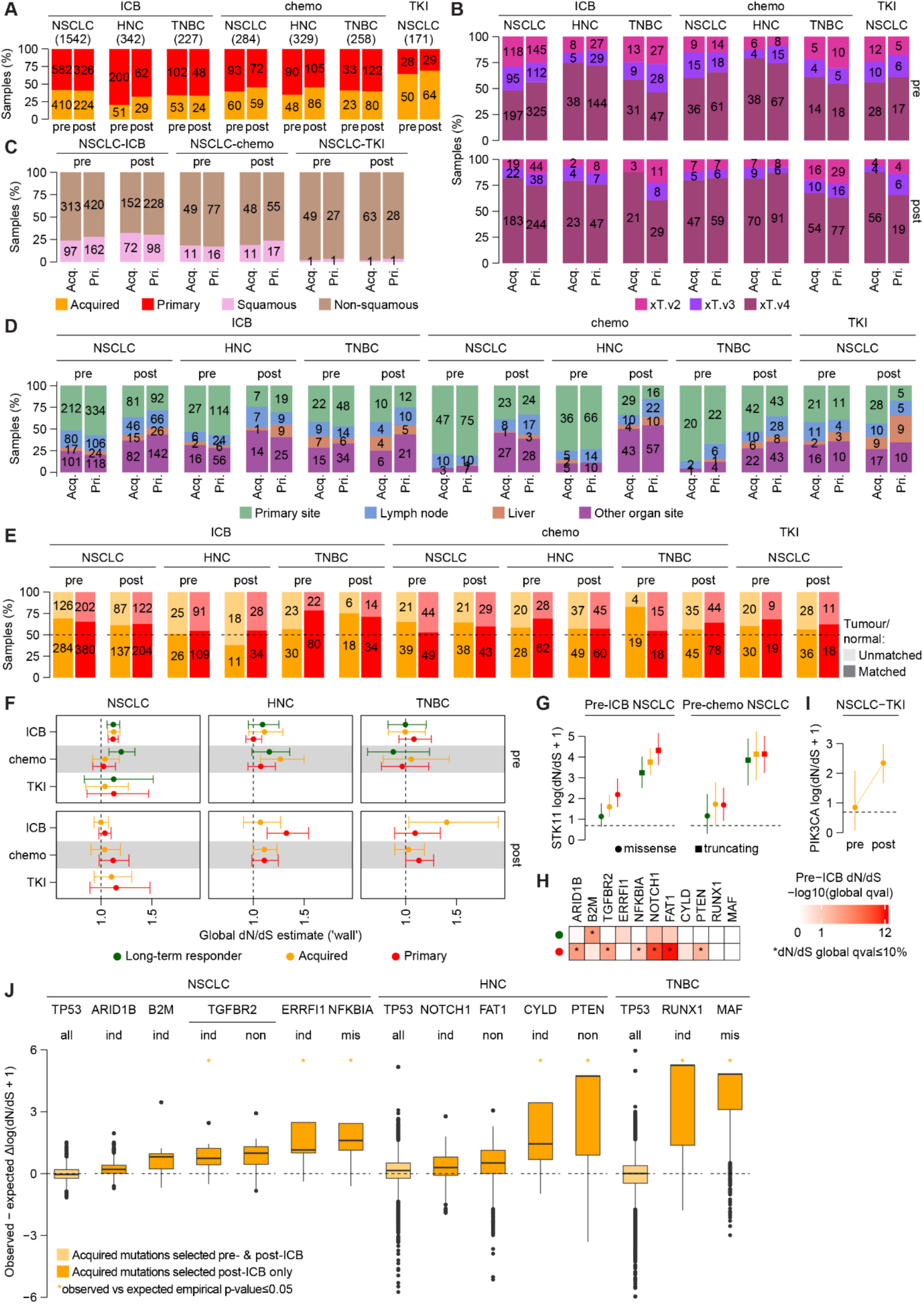
Mutation selection analysis in pre/post-treated acquired vs. primary resistant tumours. Proportion of pre-/post-treatment cross-tissue DNA-seq samples from acquired or primary resistant patients overall **(A)**, by sequencing panel **(B)**, by NSCLC histology **(C)**, by organ site **(D)**, and by matched tumour/normal sample availability **(E)** across treatment modalities and cancer types. Number of samples in each tumour group is reported. **F.** Global dN/dS estimates of all non-synonymous substitutions (i.e. ‘wall’) from *dndscv* runs in pre-/post-treatment long-term responder/acquired/primary resistant tumour groups across treatment modalities and cancer types. Error bars represent the 95% confidence intervals of the global dN/dS estimates. **G.** dN/dS estimates for *STK11* missense and truncating mutations in pre-ICB and pre-chemo long-term responder, acquired and primary resistant NSCLCs. Error bars indicate the 95% confidence intervals of the dN/dS estimates. Horizontal dashed lines designate dN/dS = 1 (i.e. neutral selection). **H.** Heatmap of dN/dS global FDR (i.e. ‘qglobal’) for acquired resistance genes in pre-ICB long-term responder and primary resistant NSCLCs (*ARID1B*, *B2M*, *TGFBR2*, *ERRFI1*, *NFKBIA*), HNCs (*NOTCH1*, *FAT1*, *CYLD*, *PTEN*) and TNBCs (*RUNX1* and *MAF*). **I.** dN/dS estimates for *PIK3CA* missense mutations in pre– and post-TKI acquired resistant NSCLCs. Error bars indicate the 95% confidence intervals of the dN/dS estimates. Horizontal dashed lines designate dN/dS = 1 (i.e. neutral selection). **J.** Distribution of the difference between the observed and expected dN/dS estimates (ΔdN/dS) for *TP53* mutations (used as negative control) and acquired resistance mutations positively selected post-ICB only. The mutation type driving selection is reported under the gene name. Each dot represents one of 10,000 iterations of the random shuffling process performed to get an expected dN/dS estimate for the corresponding gene mutation (Methods). For *TP53*, all mutation types (missense, nonsense, indels) are pooled together for a total of 30,000 data points in each cancer type. ΔdN/dS values >0 indicate higher-than-expected dN/dS estimates. Asterisks indicate mutations whose observed dN/dS is significantly higher than expected (empirical p-values). NSCLC = non-small cell lung cancer, HNC = head and neck squamous cell cancer, TNBC = triple-negative breast cancer, ICB = immune checkpoint blockade, chemo = chemotherapy, TKI = tyrosine kinase inhibitors, mis = missense, non = nonsense, ind = indels.

To understand selection dynamics before and after treatment, we contrasted dN/dS estimates of mutations positively selected post-ICB with corresponding pre-ICB estimates for both acquired and primary resistant tumours (Figure 4B). As expected, mutations selected for post-ICB included cancer-specific drivers that were also selected pre-ICB and in both acquired and primary resistance (Figure 4B). This included *STK11* in NSCLC, for which we confirmed stronger and specific pre-ICB selection in primary resistant tumours as compared to ICB-sensitive tumours^31^ (Supplementary Figure 4G). On the other hand, a subset of post-ICB mutations was identified in the post-ICB analysis but was not detected pre-ICB (i.e. post-only mutations). This was the case for all cancer types and even in post-primary resistance (except primary resistant TNBCs), indicating that ICB alters the genetic make-up of tumours regardless of resistance phenotype (Figure 4B).

In acquired resistant NSCLCs, we confirmed selection for *B2M* loss-of-function (LoF) mutations post-, but not pre-, ICB (Figure 4B), as previously described^8,12,13^. Interestingly, primary resistant NSCLCs and HNCs also showed selection for *B2M* mutations post-ICB, suggesting systematic immune selection affecting APM after treatment. We also identified mutations which have not been previously reported in the context of acquired resistance to ICB in NSCLC, including selection for *TGFBR2* (TGF-β signalling), *ERRFI1* (EGFR signalling), *ARID1B* (chromatin remodelling) and *NFKBIA* (NF-κB signalling) mutations post-ICB (Figure 4B). Likewise, we found evidence for *NOTCH1*, *FAT1*, *PTEN*, and *CYLD* LoF mutations in acquired resistant HNCs and *RUNX1* LoF and *MAF* missense mutations in acquired resistant TNBCs.

These post-only acquired ICB mutations were more recurrent post-ICB acquired resistance than both pre-ICB acquired resistance and, except *FAT1* in HNC, post-ICB primary resistance (Figure 4C). Further consolidating their involvement in ICB resistance, more than half of acquired ICB mutations were positively selected pre-ICB primary resistance but, except *B2M* in NSCLC^32^, not in pre-ICB long-term responders (Supplementary Figure 4H). Moreover, with the exception of *PTEN* and *NOTCH1* in HNC, these acquired ICB mutations were not selected for post-chemotherapy acquired resistance (Figure 4C), indicating treatment-specific selection. This was particularly striking for *CYLD* in HNC and *RUNX1* in TNBC, despite these two cancer types being more statistically powered in the chemotherapy cohort (Supplementary Figure 4A). Similarly, while we confirmed selection for *PIK3CA* mutations following acquired TKI resistance in NSCLC^33,34^, none of the acquired ICB mutations were selected for post-TKI acquired resistance (Figure 4D, Supplementary Figure 4I).

To further evaluate the extent to which these mutations are specific to acquired ICB resistance, we compared their observed dN/dS estimates to an expected distribution of dN/dS estimates (Figure 4A, Methods). Across all cancer types, while the observed post-ICB dN/dS estimates were as expected for mutations identified both pre– and post-acquired ICB resistance (e.g. *TP53* mutations), post-only acquired mutations had higher-than-expected dN/dS estimates overall (Figure 4E) and at the single-gene level (Supplementary Figure 4J). Taken together, these results emphasise the ICB-specificity of the identified acquired resistance mutations and suggest a role for these mutations in driving ICB resistance after initial clinical benefit.

### Acquired ICB alterations associate with immune-escaped TMEs

The interplay between cancer alterations and the immune TME plays a crucial role in determining response to ICB^35–38^. We next sought to investigate whether the observed selection of mutations post-ICB in the acquired resistance setting could be linked to ICB-induced immune escape. To address this, we characterised the phenotype of these mutations by comparing the transcriptional profiles of altered (ALT) vs. wild-type (WT) post-ICB acquired resistant tumours. We first expanded the repertoire of genomic alterations to include copy number variants (CNVs) and focused on mutations with a potential detrimental impact on the encoded protein (Methods). After expanding to CNVs, the recurrence of acquired ICB alterations remained highest in post-ICB acquired resistant tumours, relative to both pre-ICB acquired resistance and non-ICB regimens (Figure 5A, Supplementary Figure 5A). Collectively, the identified acquired ICB alterations affected 25-55% of acquired resistant tumours post-ICB, with <20% of altered samples carrying more than one acquired ICB alteration in the corresponding cancer type (Figure 5B).

**Figure 5.**
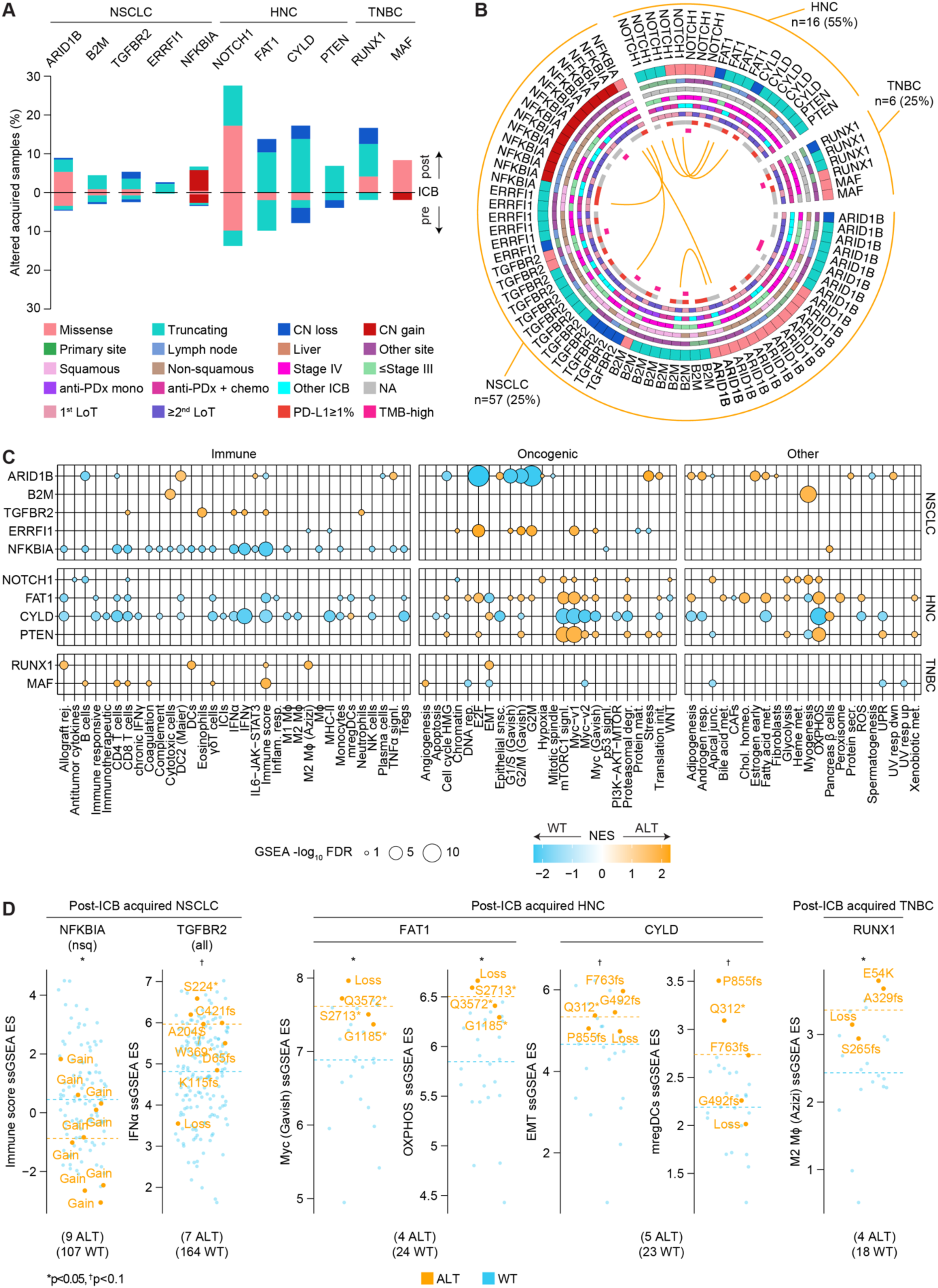
Acquired ICB alterations predict contrasted but escaped immune TMEs. **A.** Stacked barplots showing the recurrence of acquired ICB alterations in acquired resistant tumours pre– and post-ICB. Bars are annotated by alteration type for each gene. For *NFKBIA*, one sample carries multiple missense mutations and one truncating mutation and both mutation types are counted once in the barplot. The total number of samples in each tumour group (i.e. denominator) is reported in Supplementary Figure 4A. **B.** Chord diagram of post-ICB acquired resistant tumours carrying acquired ICB alterations across the three cancer types. The diagram is split by cancer type and the proportion of post-ICB acquired resistant tumours carrying acquired ICB alterations in that cancer type is reported in brackets. Each fragment represents a sample carrying the labelled acquired ICB alteration and each arc links alterations co-occurring in the same sample. The alteration type, organ site, histology, disease stage, type of ICB received, line of treatment, PD-L1 and TMB status are annotated for each sample. For *NFKBIA*, the sample labelled with missense mutation carries multiple missense mutations as well as a truncating mutation; missense is labelled as it is the mutation type driving the dN/dS signal. NA = not applicable/available and TMB-high refers to tumours with TMB ≥ 10 mutations/Mbp. **C.** GSEA NES heatmap of differentially enriched gene sets in post-ICB acquired resistant tumours carrying acquired ICB alterations (ALT) vs. WT. GSEA was performed separately for each acquired ICB alteration, focusing only on post-ICB acquired resistant tumours in the corresponding cancer type. Bubbles are proportional to-log_10_ FDR and shown only for significantly enriched gene sets (FDR≤10%). FDR was computed using Benjamini-Hochberg approach. Only gene sets differentially enriched in at least one comparison in any cancer type are shown. See Supplementary Table 6 for all GSEA results, gene set abbreviations, and number of samples used in each comparison. **D.** Distributions of ssGSEA ES of differentially enriched gene sets in ALT vs. WT post-ICB acquired resistant tumours across the three cancer types. Distributions were compared between ALT vs. WT samples using Wilcoxon rank sum test. The ES scale is divided by a factor of 1000. Dashed lines indicate median ES for ALT and WT groups. The specific gene alteration is labelled for each altered sample. For *NFKBIA*, only non-squamous NSCLC samples were used as they represent the histology underlying the GSEA enrichment. Number of ALT/WT samples is reported in brackets. NSCLC = non-small cell lung cancer, HNC = head and neck squamous cell cancer, TNBC = triple-negative breast cancer, ICB = immune checkpoint blockade, CN = copy number, anti-PDx = anti-PD(L)1, chemo = chemotherapy, mono = monotherapy, LoT = line of treatment, GSEA = gene set enrichment analysis, NES = normalised enrichment score, FDR = false discovery rate, nsq = non-squamous, ssGSEA = single sample gene set enrichment analysis, ES = enrichment score, ALT = altered, WT = wild-type.

**Supplementary Figure 5.**
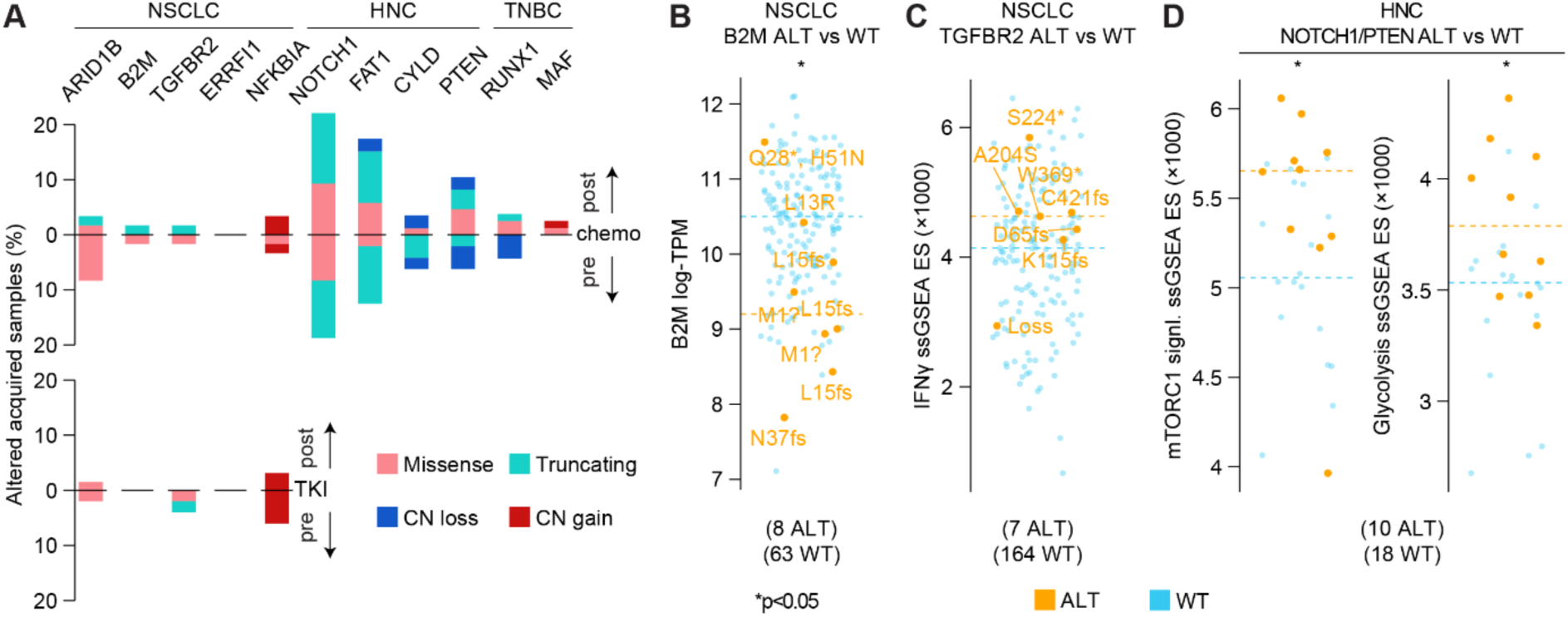
Transcriptional features of acquired ICB alterations. **A.** Stacked barplots showing the recurrence of acquired ICB alterations in acquired resistant tumours pre– and post-chemo/TKI. Bars are annotated by alteration type for each gene. The total number of samples in each tumour group (i.e. denominator) is reported in Supplementary Figure 4A. **B.** Distribution of *B2M* gene expression (TPM) in *B2M*-ALT vs. WT post-ICB acquired resistant NSCLCs. Dashed lines indicate median TPM for ALT and WT groups. Distribution of ssGSEA ES of differentially enriched gene sets in *TGFBR2*-ALT vs. WT post-ICB acquired resistant NSCLCs **(C)** and *NOTCH1*– or *PTEN*-ALT vs. WT post-ICB acquired resistant HNCs **(D)**. Dashed lines indicate median ES for ALT and WT groups. For **B**, **C**, and **D**, distributions were compared between ALT vs. WT using Wilcoxon rank sum test and the number of ALT/WT samples is reported in brackets. For **B** and **C**, the specific gene alteration is labelled for each altered sample. NSCLC = non-small cell lung cancer, HNC = head and neck squamous cell cancer, TNBC = triple-negative breast cancer, chemo = chemotherapy, TKI = tyrosine kinase inhibitors, CN = copy number, TPM = transcript-per-million, ssGSEA = single sample gene set enrichment analysis, ES = enrichment score, ALT = altered, WT = wild-type.

Comparing ALT vs. WT tumours for each acquired ICB alteration, we found two main patterns of tumour-immune associations (Figure 5C, GSEA results and number of samples used are reported in Supplementary Table 6): one group of acquired ICB alterations that associated with increased immune inflammation/infiltration (*ARID1B*, *B2M*, *TGFBR2*, *RUNX1*, *MAF*) and another group that associated with immune depletion and/or active oncogenic signalling (*ERRFI1*, *NFKBIA*, *NOTCH1*, *FAT1*, *CYLD*, *PTEN*). Importantly, these contrasted phenotypes co-existed within the same cancer type, highlighting the heterogeneity of molecular pathways linked to acquired resistance to ICB within and across cancers. For instance, in NSCLC, *B2M* LoF associated with both decreased *B2M* expression and inferred infiltration of cytotoxic cells (Figure 5C, Supplementary Figure 5B), consistent with loss of APM due to immune selection^39–42^. On the other hand, *NFKBIA* alterations, driven mainly by amplifications, associated with widespread immune depletion (Figure 5C). In HNC, tumours carrying *FAT1*, *NOTCH1*, or *PTEN* alterations showed increased proliferation (E2F/G2M/Myc), active oncogenic signalling (mTORC1/WNT), and elevated metabolism (glycolysis and oxidative phosphorylation (OXPHOS)), while *CYLD* LoF associated with predicted infiltration of immunoregulatory dendritic cells (mregDCs)^28^ and active EMT. Similarly, *RUNX1*-LoF TNBCs showed increased infiltration of dendritic cells and M2 macrophages as well as active EMT (Figure 5C).

Despite low alteration frequency and tumour sample heterogeneity, these phenotypes were generally consistent at the single-sample level (Figure 5D, Supplementary Figures 5C-D). This includes overall immune depletion in *NFKBIA*-altered non-squamous NSCLCs, contrasting with increased IFN signalling in *TGFBR2*-LoF tumours (Figure 5D, Supplementary Figure 5C). Likewise, *FAT1, NOTCH1*– or *PTEN*-LoF HNCs showed increased Myc, OXPHOS, glycolisis, and mTORC1 signalling, while *CYLD*-LoF tumours showed increased mregDCs and active EMT, and *RUNX1*-LoF TNBCs showed enrichment for M2 macrophages (Figure 5D, Supplementary Figure 5D).

Taken together, our findings suggest that ICB-induced immune responses select for acquired alterations that confer resistance to immune killing (e.g. *B2M/TGFBR2*), promote immune exclusion (e.g. *NFKBIA/RUNX1*), or enhance cancer cell-intrinsic phenotypes (e.g. *ERRFI1*/*FAT1*) to support cancer progression after initial clinical benefit.

## DISCUSSION

More than a decade after the first approvals of ICB to treat advanced-stage cancers, tumour and immune characteristics that govern the emergence of acquired resistance remain incompletely understood. In this study, we aimed to address this gap by systematically analysing patient and tumour profiles at baseline and at ICB progression, leveraging multi-modal (clinical, DNA, and RNA) and large-scale RWD. Using deep clinical and molecular metadata, we built rwICB, a real-world and multi-modal cohort of >3,500 ICB-treated patients with >2,600 pre-/post-ICB tumour samples across NSCLC, HNC and TNBC. To the best of our knowledge, rwICB constitutes the largest dataset of post-ICB resistant NSCLCs, and the first multi-modal dataset of post-ICB resistant HNCs and TNBCs. Analysing rwICB through a systematic and unbiased approach, we identified both common and cancer-specific patterns of acquired resistance, including extended survival following progression, consistent immune inflammation post-ICB, and selection for distinct acquired alterations within and across cancers.

Acquired resistance to ICB is a frequent phenomenon, estimated to occur in up to ∼70% of initial responders^3^. Yet, to this date, no standard definition of this clinical phenotype is agreed on^3,7^. While initiatives to reach a consensus definition in specific indications are ongoing^7,19,20,43^, our use of RWD and consideration of distinct cancer types across distinct treatment settings represent additional layers of complexity. Aware of these challenges, we implemented definitions that reconcile clinical precision, cohort size needs, and RWD-inherent issues, including data missingness, incompleteness, and inconsistency.

A key asset of rwICB is the availability of both rw-outcomes and long-term clinical follow-up that retrospectively allows the distinction between long-term response and acquired resistance, while also capturing primary resistance. Therefore, we used rw-outcomes conjointly with clinical follow-up to define these three response groups in rwICB. While reflecting overall expectations of outcome to ICB, including higher response rates in chemoimmunotherapy-treated patients, the proportion of acquired resistant patients in rwICB was above previous estimations (22-34% vs. ∼5-25%, overall)^3^. Besides longer patient follow-up, several factors can explain these differences. First, unlike more stringent definitions that consider only objective CR/PR as evidence of initial response^3,19^, we considered documented rwSD of at least six months as a form of response, yielding ∼20% of SD patients in our acquired ICB resistant cohorts. Second, the proportion of acquired resistant patients varied substantially across treatment and disease settings. Notably, 27-40% of chemoimmunotherapy-treated patients experienced acquired resistance, as compared to 7-27% for those treated with anti-PD(L)1 monotherapy. The latter range aligns more with estimations provided in^3^, as these were focused only on ICB monotherapy. Accordingly, the acquired resistance rate (i.e. the proportion of acquired resistant patients over all responders) at 12 months ranged between 54-58% for chemoimmunotherapy and 33-48% for ICB monotherapy in the advanced-stage setting. Therefore, our findings indicate that the incidence of acquired resistance to ICB may have been so far underestimated, emphasising the importance of treatment and disease setting when estimating the likelihood of developing acquired resistance later on.

Despite variability in incidence, acquired resistant patients consistently showed extended survival compared to primary resistant patients following ICB progression. This was the case for all cancer types and regardless of treatment or disease settings, while not observed in non-ICB modalities. We previously reported the same prognostic difference in ICB-treated NSCLC patients^8^. Here, we show that this pattern is universal and ICB-specific, indicating that an initial response to ICB is always beneficial, as its effect persists even after the onset of resistance.

Leveraging post-ICB bulk RNA-seq data and immune deconvolution, we found that acquired resistant tumours were consistently more immune-inflamed than primary resistant tumours. This immune-inflamed profile was either comparable to pre-ICB levels or exacerbated post-ICB, suggesting a sustained or enhanced immune response at the time of acquired resistance. Of note, alternative immune checkpoints, including *LAG3*^44^, were also upregulated post-ICB acquired resistance. This is in line with previous reports^16–18^ and suggests that alternative checkpoint inhibitors^45^, including bi-specific antibodies^46,47^, could reinstate an effective anti-tumour response in a subset of patients who acquired resistance to PD-(L)1/CTLA-4 inhibitors.

While the inflamed immunophenotype of acquired resistant tumours post-ICB aligns with our previous observations in a smaller cohort of matched pre/post-ICB NSCLCs^8^, we note that Ricciuti and colleagues^13^ reported significant decrease in T cell populations in acquired resistant NSCLCs post-ICB. These findings are not necessarily in contradiction with ours for several reasons. First, our use of bulk RNA-seq and the unavailability of matched pre/post-ICB samples limit our ability to detect specific immune cell changes at the single patient resolution. Second, immune TMEs can still be inflamed but T cell-excluded^38,48^. This is partly what we are observing with enriched inflammatory responses and myeloid cell signatures in post-vs. pre-ICB acquired resistance, but no significant changes in predicted T cell composition. With our larger cohort size, this lack of evidence in T cell changes is likely confounded by a mix of acquired resistant patients with either increase or decrease in T cell infiltration post-ICB, as recently reported in NSCLC^18^. In any case, these independent observations highlight the widespread inter-patient immune heterogeneity in the context of acquired resistance to ICB.

Unlike acquired resistant tumours, primary resistant tumours showed significant immune depletion from pre-to post-ICB, supporting a radically distinct immunophenotype at the onset of acquired vs. primary ICB resistance. This is also in radical contrast with the systematic immunogenic effect of chemotherapy^49,50^, which we also observe in our data regardless of resistance phenotype. Therefore, our findings provide evidence for a highly conserved and ICB-specific immunophenotype in the context of acquired resistance.

Using dN/dS to assess causal selection from pre-to post-treatment, we identified both known and novel drivers of acquired resistance. We confirmed selection for *B2M* LoF as a hallmark of acquired resistance to ICB in NSCLC^8,12,13^, while also confirming selection for *PIK3CA* mutations post-TKI acquired resistance^33,34^. Thus, these positive controls corroborate our cohort annotations and provide a solid foundation for the interpretation of our novel findings.

All acquired alterations selected for post-, but not pre-, ICB were cancer-specific, and >80% (9/11) were not selected for post-chemo/TKI, indicating high context specificity. These acquired ICB alterations affected distinct hallmarks of cancer, from dampened anti-tumour immunity to uncontrolled proliferation and dysregulated metabolism^51^. In NSCLC, *ARID1B*-LoF tumours showed increased inflammation together with decreased overall immune infiltration and cell proliferation. This aligns with a CRISPR screen where loss of chromatin remodelling components, including *ARID1B*, showed no phenotype *in vitro* but was selected for in the presence of an adaptive immune system *in vivo*^40^. Similarly, *TGFBR2* LoF associated with increased IFN signalling and immune infiltration, suggesting immune selection for the immunomodulatory role of TGF-β signalling^42^. On the other hand, *NFKBIA* alterations associated with significant immune depletion, suggesting immune exclusion via NF-κB pathway alterations. While not as immune-excluded as *NFKBIA*-altered tumours, *ERRFI1*-LoF NSCLCs showed a hyperproliferative phenotype. ERBB Receptor Feedback Inhibitor 1, or *ERRFI1*, encodes MIG6, a negative regulator of EGFR signalling^52,53^. In genetically engineered mouse models (GEMMs), loss of MIG6 accelerated both the formation and the progression of lung tumours^54^, thus supporting the highly proliferative phenotype we are observing in the context of acquired resistance to ICB. Of note, evidence for both chromatin remodelling and EGFR pathway alterations was previously found in the context of acquired ICB resistance in NSCLC, though via *SMARCA4* and *ERBB2* alterations, respectively^13^.

In HNC, besides active immunomodulatory oncogenic signalling, there was also strong evidence for a hypermetabolic phenotype in *NOTCH1*-, *FAT1*-, or *PTEN*-LoF tumours, with increased OXPHOS and glycolysis. Metabolic reprogramming in cancer cells can impact the tumour-immune cross-talk^55^ and has been implicated in acquired ICB resistance in melanoma^56^ and NSCLC^8^. Therefore, therapeutic strategies targeting altered metabolism^57^ may reverse this phenotype and restore effective anti-tumour immunity in *NOTCH1*/*FAT1*/*PTEN*-driven acquired resistant HNCs. While lacking a hypermetabolic phenotype, *CYLD*-LoF HNCs showed enrichment for mregDCs^28^ and active EMT. Cylindromatosis, or *CYLD*, is a HNC tumour suppressor that negatively regulates NF-κB signalling^58,59^ and inhibits EMT-mediated invasiveness^60,61^. Moreover, loss of *CYLD* was found positively selected in the context of an adaptive immune response in an *in vivo* CRISPR screen, but showed no phenotype *in vitro*^40^, suggesting that the tumorigenic effect of *CYLD* LoF is likely due to its immunomodulatory role. Similarly, *RUNX1*-LoF TNBCs showed increased infiltration of M2 macrophages and active EMT. The association between *RUNX1* alterations and macrophage infiltration in breast cancer was previously reported in the untreated setting^62^. Therefore, our findings point towards an involvement of the myeloid compartment in driving acquired resistance to ICB via cancer driver gene alterations. With an increasing number of clinical trials testing the use of myeloid-targeting therapies^63,64^, these alterations can serve as biomarkers to both monitor the emergence of resistance and select patients most susceptible to benefit from these next-generation immunotherapies.

Overall, our phenotypic characterisation of acquired ICB alterations nuances the generally immune-inflamed profile of acquired vs. primary resistant tumours and supports the co-existence of distinct immunophenotypes driven by specific cancer alterations. This indicates that genetic alterations are a major driver of heterogeneity in acquired resistance to ICB. We note that, with the exception of TNBC, primary resistant tumours also showed selection for *de novo* alterations post-ICB, suggesting that ICB profoundly alters the evolutionary trajectory and the clonal composition of all resistant tumours. A question arises as to why ICB induces further immune depletion and genetic remodelling post-primary resistance. A hypothesis could be that, although some determinants of resistance were already present pre-treatment (e.g. low TMB, low immune infiltration, and/or resistance mutations), additional escape mechanisms, including enhanced immunomodulatory oncogenic signalling^35,37^, may be required to sustain tumour progression with increasing ICB-induced immune pressure. Therefore, therapeutic strategies aimed at rechallenging the immune system may not benefit cases with primary ICB resistance, who may instead benefit from therapies targeting diverse oncogenic signalling pathways, including Myc signalling^65^.

Despite its systematic and comprehensive nature, our study presents several limitations, some of which were previously discussed. First, the unavailability of matched pre– and post-treatment tumour biopsies complicates the interpretation of our findings at the single patient level. Nevertheless, we addressed this limitation analytically by contrasting all our findings with 1) pre-ICB acquired vs. primary resistance, 2) post-ICB primary resistance, and 3) non-ICB regimens. In any case, both clinical and pre-clinical studies leveraging GEMMs, humanised models, single-cell and CRISPR technologies^66^ are needed to validate our findings and provide mechanistic insights into the role of each identified driver. Second, the use of targeted DNA-seq limits the scope of identifiable drivers of resistance. While the depth of coverage (>500x) in our samples likely contributed to the identification of rare events within the covered genes, future studies using whole genome/exome sequencing will complement the repertoire of alterations underlying ICB resistance. Finally, clinically refined resistance definitions accounting for oligo-progressive patterns, localised vs. spread progression, pseudo-progression, or treatment discontinuation due to toxicity^3,7^ were precluded by the limited granularity of clinical annotations in rwICB. While our definitions effectively capture the bulk of each response group, future investigations could address whether distinct molecular features define these refined clinical subgroups.

Altogether, we provide a comprehensive overview of universal and cancer-specific ICB resistance patterns, with potential for guiding treatment strategies post-progression to optimise outcomes. With an increasing awareness of the potential behind RWD^67^ and the establishment of ICB as part of routine care, we anticipate that our understanding of primary and acquired ICB resistance will rapidly expand, ultimately benefiting a larger portion of unresponsive or relapsing patients.

## METHODS

### Patient cohort building and response annotation

This study did not involve human subjects research. Data were de-identified in accordance with HIPAA. Tempus AI, Inc., has been granted an institutional review board exemption (Advarra Pro00072742) permitting the use of de-identified clinical, molecular, and multimodal data to derive or capture results, insights, or discoveries.

A sample of de-identified records from cancer patients aged 18 years old or above with early– or advanced-stage disease was selected from the multi-modal (clinical and DNA-/RNA-seq data) US-based Tempus AI real-world clinicogenomic database. Patients were enrolled in diverse oncology practices across the US, including both academic medical centres and community practices. Clinical data (patient and disease characteristics, treatment journey, and response/progression outcomes) were abstracted from structured and unstructured longitudinal physician progress notes, pathology reports, and radiology reports, using Tempus AI’s proprietary abstraction application following data ingestion and cleaning.

Starting from a total of 22,952 selected patients across ten cancer types (non-small cell lung cancer, breast cancer, head and neck cancer, bladder cancer, gastric cancer, small cell lung cancer, colorectal cancer, biliary tract/hepatocellular carcinoma, prostate cancer or ovarian cancer) and with follow-up until August 2023, we focused on those who:

- Received at least one line of treatment with complete treatment information:
  ○ care plan database identifier (ID),
  ○ name and type of medication(s) received, and
  ○ treatment start date;
- Received at least one line of ICB-containing (anti-PD-1, anti-PD-L1, or anti-CTLA-4) regimen;
- Had at least one treatment-specific real-world response event (CR/PR/SD) and/or progression event (PD or report of recurrence/metastasis) for the first line of ICB received, and these events must have been:
  ○ reported >14 days after treatment start to ensure minimum drug exposure, and
  ○ reported before any subsequent line of treatment;
- In case of presence of response event(s) only, then the patient must have been followed up for at least six months (180 days) after treatment start

No filtering on disease stage or on overall patient health profile was applied. These filtering steps yielded 4,394 ICB-treated patients across the ten cancer types, out of which the top three largest cancer types (NSCLC, HNC and breast cancer) represented >85% of the whole cohort (the next largest cancer type was bladder cancer (n=175), representing <4% of the cohort). We therefore considered these three cancer types for subsequent clinical and molecular analyses. For breast cancer, we further focused on TNBC, the subtype where ICB is indicated, to mitigate subtype-related biases in subsequent analyses. This yielded 2,689 NSCLC, 516 HNC, and 354 TNBC patients, for a total of 3,559 patients in the ICB cohort.

In parallel to the ICB cohort, we built a non-overlapping cohort of non-ICB-treated patients across the same three cancer types. The non-ICB cohort consisted of the rest of patients who had no record of ICB-containing regimen in their treatment journey and who had received either chemotherapy (for all three cancer types) or TKI (for NSCLC) as their first-line treatment. Applying the above filtering criteria yielded 929 NSCLC, 462 HNC, and 503 TNBC non-ICB-treated patients, for a total of 5,453 multi-modal patients in the built ICB and non-ICB cohorts. With the exception of 51/503 (10%) TNBCs from the chemotherapy cohort, all patients in the ICB and non-ICB cohorts (5,402/5,453 = 99%) received their treatment between 2016 and 2023, with a median treatment start year of 2020 in the ICB cohort, and 2019 in the non-ICB cohort.

To define response groups, we used treatment-specific response and progression events available for each patient and applied the following definitions for both the ICB and non-ICB cohorts:

- Long-term responders were defined as patients who had:
  ○ Documented CR/PR/SD, and
  ○ No report of progression, and
  ○ Follow-up ≥180 days from treatment start
- Primary resistant patients were defined as those who had:
  ○ Documented progression (PD/recurrence/metastasis) as their first reported outcome, or
  ○ Report of SD followed by documented progression <180 days of treatment start, or
  ○ Report of (pseudo-)response events (CR/PR) rapidly followed by documented progression <6 weeks (42 days) of treatment start
- Acquired resistant patients were defined as those who had:
  ○ Documented CR/PR followed by documented progression ≥42 days of treatment start, or
  ○ Documented SD followed by documented progression ≥180 days of treatment start.

In cases where the first outcome was SD, we always checked for and considered the subsequent non-SD event, if any, to implement these definitions. For primary resistant patients who had a progression event as their first reported outcome, no time restriction was applied on when progression was reported with respect to treatment start. This is to prevent introducing biases in survival analyses.

### Annotation of clinical and sample metadata

For accurate disease staging, stage information was parsed from both *primary_tumor_characterization* (reporting TNM staging) and *metastasis* (reporting metastasis events) files and ‘metastatic’ patients were grouped with Stage IV patients. In case of multiple stages reported at different timepoints for the same patient, we considered the stage reported closest to the treatment start date. Number of patients missing data for clinical features compared (gender, age, ethnicity, albumin level, smoking status, and stage) is reported in Supplementary Table 3.

Lines of treatment were defined in terms of drug exposure, with ‘1^st^ LoT’ referring to patients who had not previously received a drug-containing treatment, and ≥2^nd^ LoT referring to patients who received non-ICB drug(s) before the ICB regimen. The non-ICB drug can be part of a completely different treatment regimen, or part of a sequential ICB combination therapy.

For all patients in the built ICB and non-ICB cohorts, we excluded blood-based/non-solid biospecimens and focused only on solid tumour tissues biopsied from primary or distant organ sites. Pre-treatment samples consisted of any sample collected before treatment start date. All post-treatment samples were collected on or after treatment start date and before any subsequent LoT. When more than one pre-treatment sample was available per patient, we chose the one collected closest to treatment start date, or chose one arbitrarily in case of collection date ties. When more than one post-treatment sample was available per patient, we chose the one collected closest to the date of progression, or chose one arbitrarily in case collection date ties or absence of progression. DNA-seq samples were further restricted to those sequenced with xT.v2, xT.v3 or xT.v4 target sequencing assays (see DNA-seq Methods section). These sample-level filtering steps yielded a total of 4,100 pre-/post-treatment tumour samples available from 3,910 patients in the built cohorts. Of these 4,100 samples, 3,247 had RNA-seq data, 3,743 had DNA-seq data, and 2,890 had both RNA– and DNA-seq data.

### Hormone receptor status, tumour content, and PD-L1 quantification

For breast cancer patients, hormone receptor status was derived from a mix of curated (e.g. extracted from clinical notes) and native (EHR) data sources. Tumour content (purity) was determined by board certified pathologists prior to DNA extraction from formalin-fixed paraffin embedded (FFPE) tumour tissue and consisted of the proportion of tumour nuclei over all nucleated cells. PD-L1 protein expression status was either obtained externally or determined by Tempus clinical testing with the 22C3 anti-PD-L1 antibody (Agilent), as previously described^68^. Briefly, slides were scored by a pathologist using the tumour proportion score (TPS), calculated as the percentage of tumour cells with complete or partial membrane staining. Where multiple PD-L1 readouts were reported for a patient, we used the one closest to treatment start.

### Survival analyses

Survival analyses were performed for two endpoints: 1) five-year real-world overall survival (rwOS) calculated from treatment start, and 2) five-year real-world post-progression survival (rwPPS) calculated from progression. Dates of death were captured either from retrospectively abstracted patient medical records or from third-party data sources that come from obituary documentation that is augmented to the death master file from the Social Security Administration. The resulting death information was integrated with Tempus AI data via an encrypted token system. We excluded patients with missing last-known follow-up date (LKD) or death information (n=1/5,453) and those with follow-up or censoring discrepancies (no death event but reported death date, death event but no reported death date or death date different from LKD, n=29/5,453). To address survival overestimation in real-world clinicogenomic data, we performed risk set adjustment (RSA) using the patient’s first tumour sequencing date to account for delayed entries^21^. In other words, a patient was considered at risk (i.e. entry time) starting from the index date (treatment start for rwOS or progression date for rwPPS) and only after their tumour underwent next-generation sequencing. We further excluded patients whose sequencing was performed at or after LKD or more than five years after index date (n=979 for rwOS, n=855 for rwPPS). For rwPPS, we also excluded acquired/primary resistant patients whose progression date coincided with LKD (n=41). This yielded a total of 4,444 long-term responder, acquired and primary resistant patients for rwOS and 3,773 acquired and primary resistant patients for rwPPS.

We ran *survfit* function from ‘survival’ R package version 3.3.1 to fit the following RSA model and generate Kaplan-Meier curves with survival medians and associated 95% confidence intervals (based on the cumulative hazard or log(survival)):

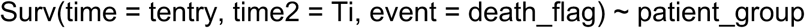

where ‘tentry’ is the delayed entry time (i.e. index date if sequencing was performed before or on index date, or date of sequencing if sequencing was performed after index date), ‘Ti’ is LKD, ‘death_flag’ is the death event variable (1 for death, 0 for censored), and ‘patient_group’ is the patient response label (long-term responder, acquired or primary resistant). If LKD was five years or more after index date, LKD was capped to five years and the patient was censored (i.e. death_flag was updated to 0).

We then ran *coxph* function from the same package on the above RSA model to fit a Cox proportional hazards regression model, compute hazard ratios and associated 95% confidence intervals, and perform likelihood ratio tests comparing all three patient groups (rwOS) or acquired vs. primary resistant patients (rwPPS).

All survival analyses were run separately for each treatment modality and cancer type and without adjusting for any co-variate. Instead, we stratified both rwOS and rwPPS analyses by disease stage (Stage IV vs. Stage I-III), LoT (1^st^ vs. ≥2^nd^), and (non-)ICB treatment modalities within each cancer type.

### RNA sequencing and data processing

RNA-seq was performed on FFPE tumour samples using the Tempus xR.v1 or xR.v2 exome-capture-based assays (Tempus AI, Inc., Chicago, IL). All RNA extraction, library preparation, sequencing and data processing are described in^69^. Briefly, Tempus xR assays are whole-transcriptome next-generation sequencing assays spanning a 39 Mb target region of the human genome covering a total of 19,375 genes. Sequencing was performed with Illumina HiSeq 4000 (RS.v1) or NovaSeq 6000 (RS.v2) and reads were aligned to GRCh37. Gene-level expression was quantified as both raw counts as well as gene length-, G+C content-, and library size-normalised transcripts per million (TPM). TPM values were log_2_-transformed and batch-corrected for assay version using an in-house approach^69^. All subsequent analyses used either raw count data (DEA) or batch-corrected log_2_-transformed TPM data (single gene comparisons, UMAP, and ssGSEA).

### Differential expression and gene set enrichment analyses

We used DESeq2^70^ R package version 1.32.0 to run a contrasted DEA model comparing the transcriptional profiles of pre-/post-treatment acquired/primary resistant tumours within each cancer type and treatment modality separately (chemo for all three cancer types; anti-PDx mono, anti-PDx + chemo, other ICB and TKI for NSCLC; any ICB for all three cancer types in addition to 1^st^ LoT Stage IV NSCLC, ≥2^nd^ LoT Stage IV NSCLC, squamous NSCLC, and non-squamous NSCLC). We used raw count gene expression data of tumours biopsied from the primary organ site for each cancer type (lung tissue for NSCLC, head/neck tissue for HNC, breast tissue for TNBC) and ran the following model accounting for RNA-seq assay version (RS.v1/2) and NSCLC histology (squamous/non-squamous), leaving all parameters to default:

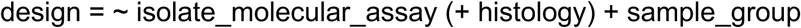

where ‘sample_group’ is a concatenation of pre-/post-treatment and acquired/primary resistant sample type (i.e. one of pre_acquired, post_acquired, pre_primary, post_primary).

DEA results were then extracted specifying the contrast of each comparison of interest and setting the significance threshold (i.e. ‘alpha’) to 0.05:

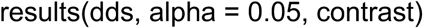

where ‘dds’ is the fitted DEA object and ‘contrast’ is one of:

- Pre acquired vs. pre primary
- Post acquired vs. post primary
- Pre acquired vs. post acquired
- Pre primary vs. post primary

The reference group of each comparison and the number of samples used are reported in Supplementary Table 4. Log_2_ fold changes (log_2_FC) and associated p-values were computed using Wald tests and p-values were adjusted for multiple tests using Benjamini-Hochberg correction.

We then used DESeq2 results to run pre-ranked GSEA using *fgseaMultilevel* function from fgsea^71^ R package version 1.20.0 with default parameters. Specifically, all 19,375 genes were ranked based on their DESeq2 log_10_(FDR) signed by the log_2_FC sign, with genes ranked on the top being the most significantly upregulated in the reference group, and the ones in the bottom being the most significantly downregulated. Genes whose DESeq2 FDR = NA were excluded. For each gene set, *fgseaMultilevel* walks through the ranked gene list and computes a running sum statistic-based enrichment score with an associated p-value using the adaptive multilevel splitting Monte Carlo approach. Normalised enrichment scores were computed to account for differences in gene set sizes and were used in all comparisons. For each comparison, p-values were adjusted for multiple testing across gene sets using Benjamini-Hochberg correction. The following gene sets were considered for all comparisons and no filter on the gene set size was applied:

- 50 Hallmark gene sets from MSigDB^25^;
- 19 cancer-specific immune cell signatures from ConsensusTME^23^ (TCGA-LUAD for NSCLC, TCGA-HNSCC for HNC, and TCGA-BRCA for TNBC);
- 28 literature-compiled immuno/oncology-focused gene sets: 4 dendritic cell signatures (DC1, DC2, mregDC, cDCmaturation) from^28^, 2 macrophage signatures (M1 and M2 macrophages) from^27^, 9 TME signatures (MHCI, MHCII, coactivation molecules, antitumor cytokines, protumor cytokines, checkpoint inhibition, MDSCs, CAFs, Matrix) from^29^, and 10 tumour meta-program signatures (MP1 Cell Cycle – G2/M, MP2 Cell Cycle – G1/S, MP3 Cell Cylce HMG-rich, MP4 Chromatin, MP5 Stress, MP8 Proteasomal degradation, MP10 Protein maturation, MP11 Translation initiation, MP19 Epithelial Senescence, MP20 MYC) from^26^.

For each comparison, GSEA was run together on Hallmark and ConsensusTME gene sets, and run separately on the literature-compiled gene sets. Gene sets with GSEA FDR≤0.1 were considered significantly differentially enriched.

### Single sample gene set enrichment analysis

We used *gsva* function from GSVA^72^ R package version 1.42.0 to run ssGSEA^24^ for all gene sets on the batch-corrected and log_2_-transformed TPM data of all RNA-seq tumour samples. All parameters were set to default except for ssgsea.norm which was set to FALSE (i.e. run without normalisation) to allow for absolute computation of enrichment scores (ES). Distributions of ssGSEA ES of specific gene sets were compared between tumour groups of interest using Wilcoxon rank sum test.

### DNA sequencing and data processing

DNA-seq was performed on FFPE tumour samples and was available for a total of 3,981 FFPE samples from the built ICB and non-ICB cohorts. Sequencing was performed using Tempus xT.v1-4 targeted panel (3,930/3,981=98.7% samples, 596-648 genes), xO targeted panel (18/3,981=0.5% samples, 1,714 genes) or xE.v1-2 whole exome sequencing (33/3,981=0.8% samples) assays (Tempus AI, Inc., Chicago, IL). We focused on samples sequenced with xT.v2, xT.v3, or xT.v4 assays (3,743/3,981=94% DNA-sequenced FFPE tumour samples) to maximise both the number of samples and commonly targeted genes across assays (n=595 commonly covered cancer-related genes across xTv2-4). A matched normal sample (i.e. germline control) was available for 2,352/3,743 samples (63%) and consisted of 200 μl blood or 650 μl saliva. Sequencing with xT assays was performed to achieve an average on-target depth of 500x. All DNA extraction, library preparation, sequencing and data processing, including calling of somatic mutations and CNVs (for both normal-matched and tumour-only samples), classification of mutation functional impact, calculation of TMB, detection of MSI, and HLA typing, is described in^73,74^. A total of 235,311 short variants (single nucleotide variants and indels) aligned to GRCh37 were detected across the 3,743 samples.

Given sensitivity of dN/dS analysis to germline contamination (see *dndscv* Methods section), and since a germline control was not always available for DNA-seq, we further cross-checked all mutation calls against germline variant databases (dbSNP version 154 and gnomAD version 2.1.1) and filtered out any germline variant with an allele frequency ≥0.1%. Out of the 235,311 mutations, 55,173 (23%) were filtered out, yielding a total of 180,138 mutations in the final mutation dataset, with a median mean coverage at mutated sites of 563x across samples (90% of samples with mean coverage ≥200x, 99% with mean coverage ≥100x).

Two versions of the mutation dataset were used in subsequent analyses: 1) an unfiltered version with all 180,138 mutations containing synonymous mutations and without filtering applied on variant allele frequency or mutation functional impact (version used for *dndscv* analysis), and 2) a filtered version with 67,759 non-synonymous and potentially more biologically relevant mutations after applying the filtering steps described in^73,74^ (version used for the functional characterisation of acquired ICB alterations).

### Analysis of mutation selection with *dndscv*

Using the unfiltered mutation dataset containing both synonymous and non-synonymous mutations (see DNA-seq Methods section), we ran *dndscv*^30^ (https://github.com/im3sanger/dndscv) version 0.0.1.0 in each of pre-/post-treatment acquired/primary resistant and pre-treatment long-term responder tumour groups separately within each cancer type and for each of ICB, chemotherapy, and TKI treatment modalities, for a total of 35 *dndscv* runs. Because this analysis relies on cancer mutations, we used all tumour samples available in our annotated dataset, including those biopsied from non-primary (metastatic) organ sites, to increase statistical power.

We used the hg19 reference coding DNA sequence (refCDS) object downloaded from https://github.com/im3sanger/dndscv/blob/master/data/refcds_hg19.rda. Of the 595 commonly covered genes across xT.v2-4 assays, 592 genes were present in hg19 refCDS (i.e. all but *H19*, *HLA-DPB2*, and *HLA-DRB6*). For *CDKN2A*, *dndscv* considers two isoforms separately: CDKN2A.p16INK4a and CDKN2A.p14arf. We used the following parameter settings for all *dndscv* runs: max_muts_per_gene_per_sample = 10 (i.e. the maximum number of mutations per gene per sample), outmats = T, constrain_wnon_wspl = T (i.e. constraining nonsense and essential splice site dN/dS estimates for increased power), gene_list = common_panel_genes (i.e. all 592 genes commonly covered by the three DNA-seq assays).

An important requirement for *dndscv* is the exclusion of all germline variants from the mutation dataset, as such variants typically bias dN/dS estimates towards lower values (<<1)^30^. After cross-checking all mutation calls against germline variant databases as previously described, we checked that the global dN/dS estimates across all 592 genes (‘wall’) were close to or slightly above neutral selection (≥1) for all *dndscv* runs (Supplementary Figure 4F), as expected.

For each gene and for each mutation type (missense, nonsense, indels), *dndscv* computes a maximum-likelihood estimate (MLE) of the dN/dS ratio and a corresponding p-value from the Likelihood-Ratio test, in addition to a global p-value integrating all mutation types. All p-values are adjusted for multiple testing using Benjamini-Hochberg correction (FDR). A gene was considered under significant positive selection if at least one mutation type affecting the gene (missense/nonsense/indel) was under significant positive selection, and if the global selection FDR was below significance threshold. More specifically:

- (wmis_cv > 1 and qmis_cv ≤ 0.1) or (wnon_cv > 1 and qnon_cv ≤ 0.1) or (wind_cv > 1 and qind_cv ≤ 0.1), and
- qglobal_cv ≤ 0.1

where wmis_cv, wnon_cv, and wind_cv are the dN/dS MLE estimates for missense, nonsense (truncating), and indel mutations respectively, qmis_cv/qnon_cv/qind_cv are the corresponding mutation-level selection FDRs, and qglobal_cv is the global selection FDR integrating all mutation types. Positively selected genes are represented by their mutation-level log-transformed dN/dS MLE from the corresponding *dndscv* run in Figure 4B.

Selection for *NFKBIA* mutations in post-ICB acquired NSCLCs was driven by missense mutations in two samples: one sample that had one missense mutation, and another sample that had six missense and one nonsense mutation. We checked that the latter sample was not hypermutated (TMB<10 mutations/Mbp and MSI-stable), and all seven mutations affecting *NFKBIA* in that sample had a read depth ranging from 1,361-1,630x and a variant allele frequency ranging from 8-15% for a tumour purity of 20%. Restricting the *dndscv* analysis to a maximum of 3 mutations per gene per sample (i.e. the default) in post-ICB acquired NSCLCs yielded a missense dN/dS MLE of 5.9 for *NFKBIA* with an uncorrected missense p-value of 0.01 and an uncorrected global p-value of 0.048.

### Comparison of observed vs. expected dN/dS estimates

To evaluate the specificity of mutations selected post-(but not pre-)ICB acquired resistance, we compared their observed dN/dS estimates to an expected distribution of dN/dS estimates, contrasting with the same comparison for mutations selected both pre– and post-ICB. The expected distribution of dN/dS estimates was generated by randomly shuffling pre/post-ICB acquired/primary tumour group labels and re-running *dndscv* on the pseudo-post-ICB acquired resistant group, repeating the process 10,000 times. More specifically, for each cancer type in the ICB cohort, we:

- Focused on pre_primary, post_primary, pre_acquired and post_acquired tumour samples (i.e. 4 labels);
- Randomly shuffled the 4 labels then took the pseudo-post_acquired labelled samples (this way, we ensure that we randomly sample the same number of actual post_acquired samples at each iteration);
- Ran *dndscv* on the pseudo-post_acquired samples using the same parameter settings as previously described;
- Repeated this process 10,000 times to generate a null distribution of dN/dS estimates.

Then, for each positively selected post-ICB acquired mutation, we calculated the difference between its observed dN/dS MLE and all 10,000 expected dN/dS MLEs, obtaining ΔdN/dS distributions for mutations selected post-only or pre– and post-ICB acquired resistance, as depicted in Figure 4E. At the gene level, we assessed the significance of the difference between observed vs. expected dN/dS estimates by computing an empirical p-value assessing how often the observed dN/dS estimate was higher than expected across all iterations, i.e. p-value = n(ΔdN/dS > 0)/10,000.

### Functional characterisation of acquired ICB alterations

Focusing on post-ICB acquired resistant tumour samples with both DNA– and RNA-seq data, we ran DEA and GSEA comparing ALT vs. WT tumours for each acquired ICB alteration in each corresponding cancer type separately. To annotate ALT tumours, we integrated CNV data with the filtered mutation dataset (see DNA-seq methods) and considered only pathogenic (P), likely pathogenic (LP), or mutations with unknown significance (US), excluding all benign or likely benign mutations. A gene was considered altered (ALT) depending on its cancer driver role as annotated in^75^ or on the mutation type driving dN/dS selection (i.e. missense or truncating/indel mutations):

- If the gene was a tumour suppressor gene (TSG), then only loss-of-function (LoF) alterations were considered;
- If the gene was an oncogene (OG), then only gain-of-function (GoF) alterations were considered;
- If the gene had no clearly defined cancer driver role (TSG/OG), then LoF alterations were considered if the dN/dS selection was driven by truncating/indel mutations, otherwise any alteration (P/LP/US mutation or CN loss/gain) was considered for the gene.

Any CN loss, stop gain, frameshift, or start loss mutation were considered as LoF alterations, while CN gains were considered as GoF alterations. Missense mutations affecting TSGs were considered LoF alterations, and those affecting OGs were considered GoF alterations. We considered LoF alterations for *ARID1B*, *B2M*, *TGFBR2*, *NOTCH1*, *FAT1*, *CYLD*, *PTEN*, and *RUNX1* (TSGs), as well as for *ERRFI1* (no clearly defined cancer driver role but dN/dS selection driven by indels). We considered GoF alterations for *MAF* (OG). Since no clear cancer driver role was reported for *NFKBIA* and since dN/dS selection was driven by missense mutations, we considered all alteration types for this gene.

We used raw count gene expression data of tumours biopsied from primary or metastatic organ sites (cross-tissue sites) and ran the following DESeq2 DEA model accounting for RNA-seq assay version (RS.v1/2), NSCLC histology (squamous/non-squamous), and organ site (primary site/lymph node/liver/other), leaving all parameters to default:

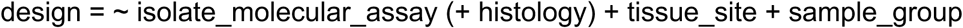

where tissue_site is the organ site and sample_group is one of ALT or WT. DEA results were extracted setting ‘alpha’ parameter to 0.05. We then ran pre-ranked GSEA using DEA results to find differentially enriched gene sets between ALT vs. WT tumours, using Hallmark, ConsensusTME, and literature-compiled gene sets as previously described.

### Statistical analyses

All analyses were performed and plots produced using R programming language version 4.1.0. Statistical tests used in each analysis/comparison are specified in the corresponding figure legends or Methods section. All p-values were adjusted using Benjamini-Hochberg correction where applicable.

## AUTHOR DECLARATION

M.R.K, R.R., Je.D., S.A.H., D.P., R.S., A.C.S., S.K. and M.L.M are AstraZeneca employees and shareholders. S.C.P is a Boehringer Ingelheim employee and shareholder. Z.K.A. is a Quotient Therapeutics employee and shareholder. B.S. is a Biorelate Ltd employee and shareholder. K.B. and Jo.D. are Tempus AI employees and shareholders. A.J.S. reports consulting/advising role to J&J, KSQ therapeutics, BMS, Merck, Enara Bio, Perceptive Advisors, Oppenheimer and Co, Umoja Biopharma, Legend Biotech, Iovance Biotherapeutics, Prelude Therapeutics, Immunocore, Lyell Immunopharma, Amgen and Heat Biologics. Research funding: GSK (Inst), PACT pharma (Inst), Iovance Biotherapeutics (Inst), Achilles therapeutics (Inst), Merck (Inst), BMS (Inst), Harpoon Therapeutics (Inst) and Amgen (Inst). I.M. is a cofounder, shareholder, and consultant for Quotient Therapeutics.

## AUTHOR CONTRIBUTIONS

M.R.K., A.J.S, and M.L.M. conceptualised the study. M.R.K. and M.L.M. designed the analysis plan. M.R.K., S.C.P., Z.K.A., K.B., and Je.D. analysed the data. M.R.K, R.R., S.A.H., D.P., R.S., I.M., A.J.S, and M.L.M. interpreted the data. A.C.S., S.A.H., D.P., B.S., Jo.D., S.K. and M.L.M. managed the project. M.R.K. and M.L.M wrote the manuscript. All authors edited the manuscript.

## DATA AVAILABILITY

De-identified data used in the research were collected in a real-world health care setting and are subject to controlled access for privacy and proprietary reasons. When possible, derived data supporting the findings of this study have been made available within the paper and its Supplementary Figures/Tables. Restrictions apply to the availability of additional data, which were used under license for this study.

## ACKNOWLEDGEMENTS

We would like to thank Matthew D. Hellmann for his insightful comments on the manuscript. This study was funded by AstraZeneca.

